# Extracellular G-quadruplex and Z-DNA protect biofilms from DNase I and forms a DNAzyme with peroxidase activity

**DOI:** 10.1101/2023.05.22.541711

**Authors:** Gabriel Antonio S. Minero, Andreas Møllebjerg, Celine Thiesen, Mikkel Illemann Johansen, Nis Pedersen Jørgensen, Victoria Birkedal, Daniel Otzen, Rikke L. Meyer

## Abstract

Many bacteria form biofilms to protect themselves from predators or stressful environmental conditions. In the biofilm, bacteria are embedded in a protective extracellular matrix composed of polysaccharides, proteins and extracellular DNA (eDNA). eDNA most often arises from lysed cells, and it is the only matrix component most biofilms appear to have in common. However, little is known about the form DNA takes in the extracellular space, and how different non-canonical DNA structures such as Z-DNA or G-quadruplex formation might contribute to its function in the biofilm.

The aim of this study was to determine if non-canonical DNA structures form in eDNA-rich staphylococcal biofilms, and if these structures protect the biofilm from degradation by nucleases. We grew *Staphylococcus epidermidis* biofilms in laboratory media amended with hemin and NaCl to stabilize secondary DNA structures and visualized their location by immunolabelling and fluorescence microscopy. We furthermore visualized the macroscopic biofilm structure by optical coherence tomography. We developed assays to quantify degradation of Z-DNA and G-quadruplex DNA oligos by different nucleases, and subsequently investigated how these enzymes affected eDNA in the biofilms.

Z-DNA and G-quadruplex DNA were abundant in the biofilm matrix, and were often present in a web-like structure in biofilms grown *in vitro* and *in vivo* using a murine implant-associated osteomyelitis model. *In vitro*, the structures did not form in the absence of NaCl or mechanical shaking during biofilm growth, or in bacterial strains deficient in eDNA or exopolysaccharide production. We thus infer that eDNA and polysaccharides interact, leading to non-canonical DNA structures under mechanical stress when stabilized by salt, and we confirmed that G-quadruplex DNA and Z-DNA was also present in biofilms from infected implants. Mammalian DNase I lacked activity against Z-DNA and G-quadruplex DNA, while Micrococcal nuclease could degrade G-quadruplex DNA and S1 Aspergillus nuclease could degrade Z-DNA. Micrococcal nuclease, which originates from *Staphylococcus aureus*, may thus be key for dispersal of biofilm in staphylococci. In addition to its structural role, we show for the first time that the eDNA in biofilms forms a DNAzyme with peroxidase-like activity in the presence of hemin. While peroxidases are part of host defenses against pathogens, we now show that biofilms can possess intrinsic peroxidase activity in the extracellular matrix.

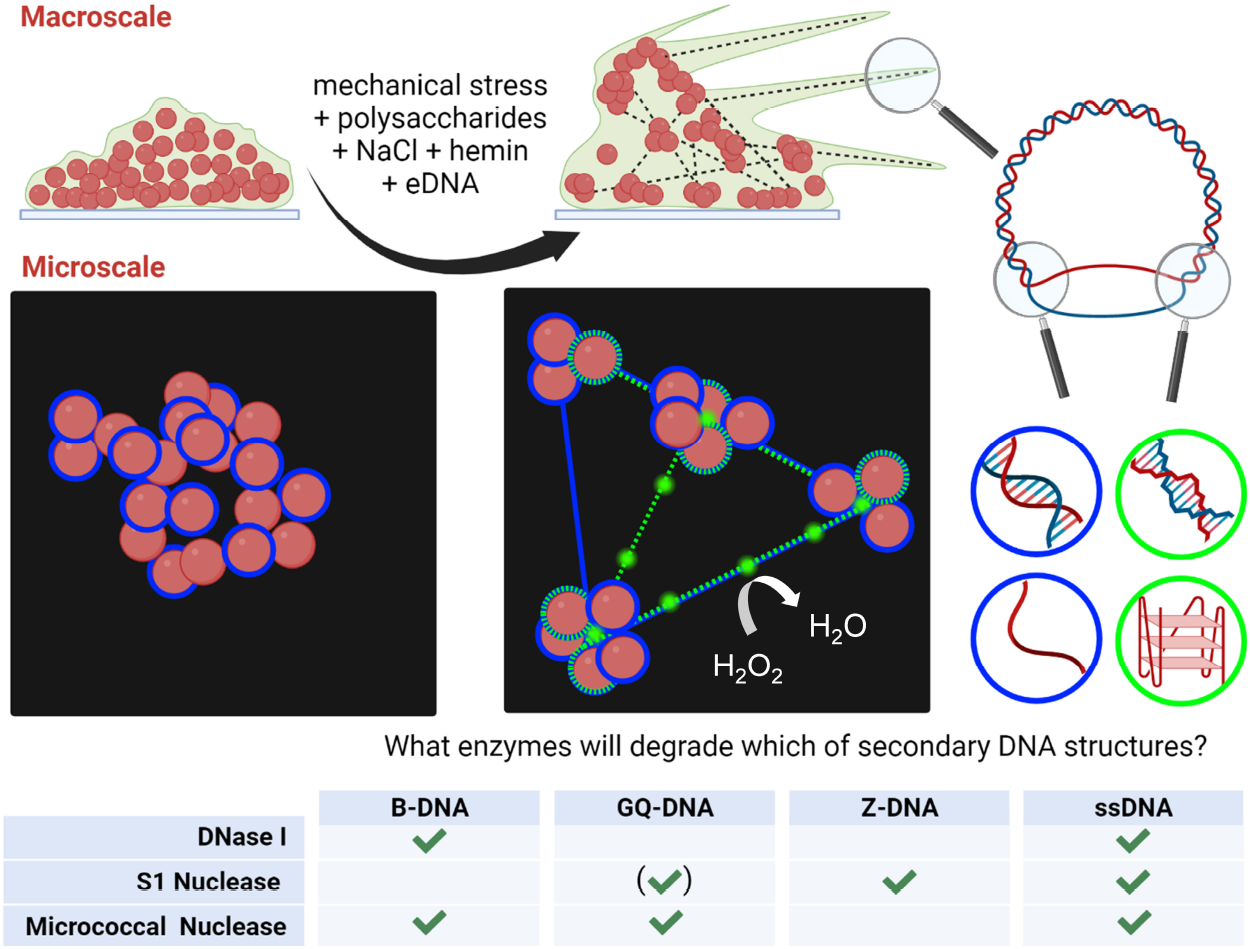

## INTRODUCTION

The most problematic bacterial infections today are those that involve biofilms: a multicellular community of bacteria encased in a shared extracellular matrix. Biofilms caused by pathogens are typically associated with implants or wound infections, and these are becoming a significant burden to the healthcare system. Biofilm formation is an important virulence factor, as it protects bacteria from the immune system and antibiotic therapy. In the US alone, the National Institutes of Health estimate that 65 % of acute and 80 % of chronic infections are associated with biofilm formation, incurring an annual economic burden of approximately 11 billion dollars (*1*).

The biofilm matrix consists of polysaccharides, proteins, and extracellular DNA (eDNA). While some bacteria produce toxins and several other virulence factors, the commensal skin bacterium *Staphylococcus epidermidis* (*S. epidermidis*) relies primarily on biofilm formation to support its existence in the human body. Thanks to its ability to form biofilm, *S. epidermidis* is a major cause of catheter-related bloodstream infections (*2*) and serious implant-associated infections such as prosthetic valve endocarditis (*3*), cardiac implantable electronic device infection (*4*) and prosthetic joint infection (*5, 6*). Due to high rates of resistance to several classes of antimicrobials, treatment failure is high in prosthetic joint infections (*6*).

Among the mechanisms for biofilm formation, *S. epidermidis* uses eDNA, poly-N-acetylglucosamine (PNAG), and a plethora of cell wall-anchored proteins that are used for attachment to the host, and other molecules (*7*). One third of clinical isolates from implant-associated infections lack the genes for polysaccharide production (*8*). In contrast, most clinical isolates encode a giant 10 kDa surface protein, Embp (*9*), which is co-regulated with release of eDNA (*7*). Thus, eDNA is important for establishment of biofilm infections by *S. epidermidis* (*10-12*) as it is in many other bacterial species (*13, 14*).

Efforts to disperse biofilms or facilitate the penetration of antibiotics (*15*) have long focused on eDNA as a prime target for enzymatic degradation of the biofilm matrix. Frustratingly, mammalian DNase I treatment only prevents formation of biofilms, while it is ineffective at breaking up mature biofilms (*14*) or biofilms obtained in media with high ionic strength (*16*). Until recently, it was assumed that eDNA exists primarily in its canonical right-handed B-form. However, Buzzo *et al*. reported on formation of the left-handed Z-DNA in various species biofilms *in vitro* as well as *in vivo* (*17*). Moreover, Seviour *et al*. identified both G-quadruplex (GQ-) DNA and RNA in *Pseudomonas aeruginosa* biofilms (*18, 19*). These non-canonical DNA structures are resistant to degradation by DNase I (*20*), and thus may explain why this enzyme fails to disperse biofilms.

In the cell, Z-DNA is found in supercoiled bacterial genomes or plasmids. Under physiological conditions, Z-DNA forms in (GC)_n_ sequences of n ≥8, while shorter (GC)_n_ sequences are in dynamic equilibrium between B- and Z-DNA conformations (*21*). *In vitro*, Z-DNA formation from linear non-supercoiled DNA oligos requires high salinity (up to 4.5 M NaCl) (*22*), polycationic molecules such as polyamines (e.g. spermine) (*23*), or cationic metalloporphyrins (*24, 25*). Furthermore, different chemical modifications of nucleobases such as methylation or protonation of cytosine (*26, 27*), or bromination or oxidation of guanine (*28*) can promote formation/stabilization of Z-DNA. It remains unknown which mechanisms control formation of Z-DNA in the extracellular environment, but Buzzo *et al*. suggested that Z-DNA predominates in Holliday junction DNA structures stabilized by DNABII proteins wherein the double-stranded DNA is wrapped or twisted around DNABII proteins (*17*). In such DNA protein complexes, the ends of DNA could be locked in a high energy state similar to a supercoiled state conducive for Z-DNA formation.

G-quadruplexes (GQ) are other secondary structures found in G-rich sequences and in supercoiled bacterial genomes under physiological conditions (*29*). Typically, three consecutive G-tetrads separated by loops of 1-7 nucleotides are required to form intramolecular GQ (*30*). Intermolecular GQs may fold from two or four single strands of DNA or RNA (*31*), as well as higher-order intermolecular GQ-wires can form under crowding conditions (*32*) or simply in the presence of 10 mM Mg divalent cations (*33*). Several environmental conditions affect GQ formation. K^+^ and Na^+^ stabilize GQ, while Cs^+^ or Li^+^ do not favor GQ formation (*34*). Simulations also suggest possible stabilization of GQ by heme, which binds to GQ with high affinity (*35*). Finally, oxidation of guanines can also affect GQ conformation and stability (*36*).

We hypothesize that the availability of specific extracellular polymeric substances, cations, metalloporphyrins, oxidizing conditions, and exposure to mechanical stress that leads to bending DNA to a state similar to supercoiled state could promote formation of Z-DNA and G-quadruplex structures after genomic DNA is released to the extracellular space in biofilms. These conditions are present in *S. epidermidis’* natural environment on the skin, in wound infections and presumably adjacent to implants located in the arteries, bones and the subcutaneous space. Furthermore, we hypothesize that non-mammalian nucleases can degrade these structures and may be used for bacteria-driven biofilm dispersal while biofilms remain protected from host nucleases.

Accordingly, the first aim of this study was to determine if non-canonical DNA structures exist in the matrix of *S. epidermidis* biofilms, and to identify environmental and biological factors that affect formation of these structures. The second aim was to determine if secondary structures are protected from degradation by mammalian DNase I and bacteria-derived nucleases. We use immunolabelling and confocal microscopy to visualize various DNA secondary structures at the μm scale in biofilms formed *in vitro* and *in vivo*, and optical coherence tomography to observe changes in the matrix structure on the μm-mm scale. We furthermore evaluate activity of three nucleases (DNase I, S1 nuclease and Micrococcal nuclease) against various secondary structures of DNA oligos, and subsequently assess how the properties of these nucleases affect their ability to degrade eDNA in biofilms. Moreover, for the first time we demonstrate extracellular peroxidase activity in biofilms.

## RESULTS

### *S. epidermidis* biofilm form a web-like extracellular matrix in the presence of hemin and NaCl

We employed optical coherence tomography (OCT) to visualize biofilms at microscale (*37*). Here we show that *S. epidermidis* 1457 formed a web-like matrix extending more than 1 mm from the underlying substrate and attaching to the sides of the microwell (Figure 1A-B) in tryptic soy broth (TSB) amended with 200 mM NaCl (TSB-NaCl), resulting in a total concentration of 285 mM NaCl. Considering *S. epidermidis*’ ability to form biofilm at elevated salinity, we hypothesize that these structures are similar to the “streamers” reported for *S. epidermidis* grown under lateral flow (*38*). Streamers are primarily made from polysaccharides and eDNA and extend hundreds of μm from the biofilm when subjected to the mechanical stress of liquid flow. In our system, the 150 rpm rotation during incubation also creates mechanical stress as the liquid swirls up the sides of the well. To investigate if eDNA and poly-N-acetyl glucosamine (PNAG) were responsible for the web-like biofilm matrix observed here, we visualized biofilms formed by mutant strains lacking the ability to produce eDNA (1457 *ΔatlE*) (Figure 1C) or polysaccharides (1457 *ΔicaADBC*) (Figure 1D). Neither of these strains produced the web-like biofilm matrix, suggesting that eDNA and PNAG are both essential components. We therefore conclude that the web-like strings of the biofilm matrix are similar to the biofilm streamers observed in lateral flow systems.

**Figure 1.**
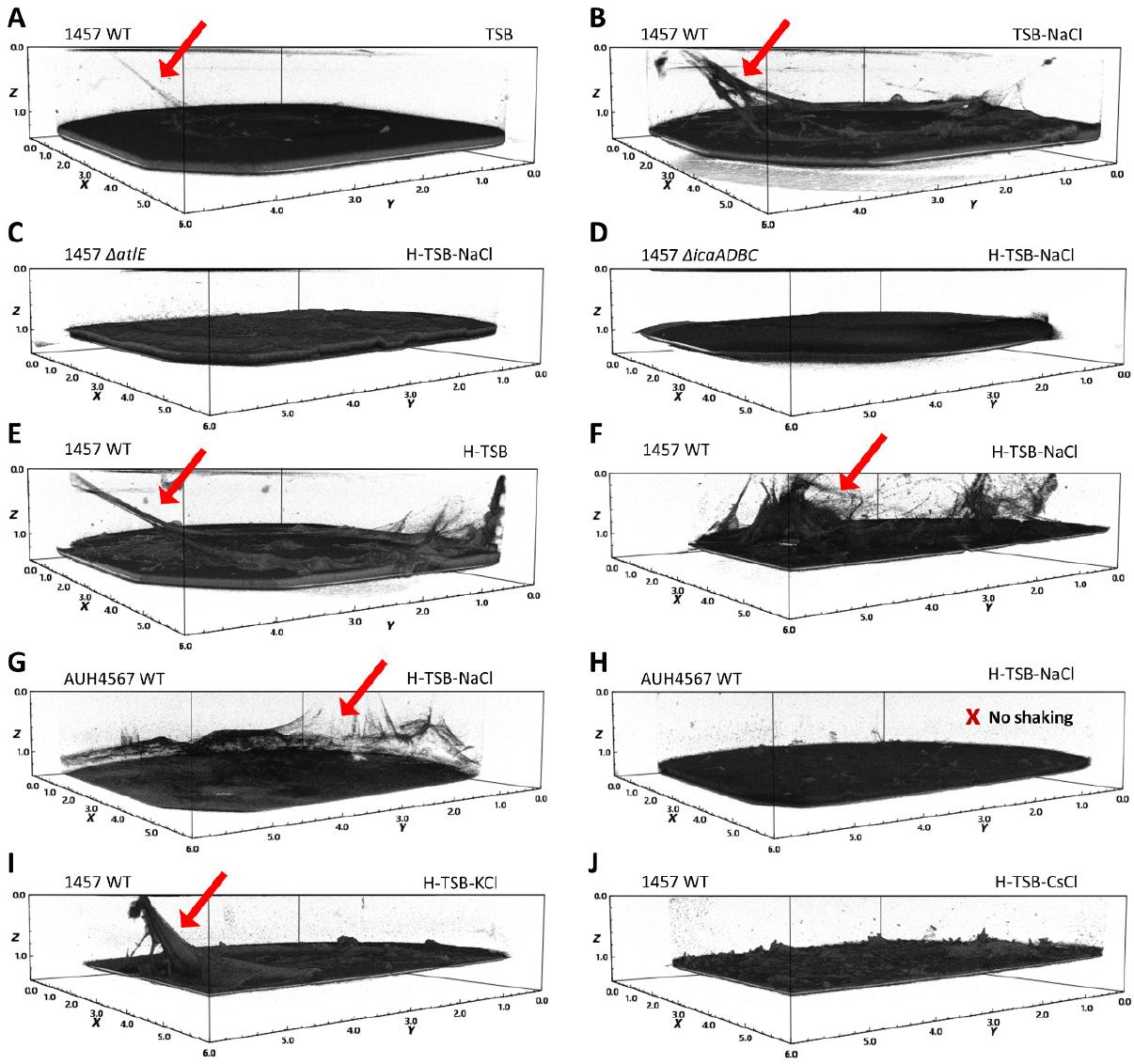
*S. epidermidis* biofilms exposed to mechanical stress produce streamers made from polysaccharides and eDNA. Formation of the streamers relies on the presence of NaCl, KCl and hemin. *S. epidermidis* biofilms were grown for 3 days in TSB with different supplements at 150 rpm rotation or without shaking (H). One biological replicate out of three is shown for (A) the 1457 WT in TSB, (B) the 1457 WT in TSB-NaCl, (C) the 1457 eDNA-deficient mutant Δ*atlE* in the H-TSB-NaCl, (D) the 1457 polysaccharide-deficient mutant Δ*icaADBC* in the H-TSB-NaCl, (E) the 1457 WT in H-TSB, (F) the 1457 WT in H-TSB-KCl, (G)-(H) the AUH4567 WT in the H-TSB-NaCl, (I) the 1457 WT in H-TSB-KCl, (J) the 1457 WT in H-TSB-CsCl. Dimensions are 6 × 6 × 1 mm. Arrows indicate examples of biofilm streamers.

We hypothesize that the mechanical stress that led to streamer formation will also promote formation of non-canonical secondary structures of eDNA due to bending of the DNA (*39*), which resembles DNA coiling (*21, 29*). Non-canonical structures may be advantageous for biofilms under high mechanical stress, as some structures are more mechanically robust (*39*). To investigate if streamers might involve non-canonical DNA structures, we explored the biofilm formation under environmental conditions that stabilize these structures. Metalloporphyrins can stabilize certain DNA structures (*25, 35, 40*), and are available to pathogens in the form of heme or hemin in the bloodstream. Hemin is known to interact with GQ (*41*), and being a metalloporphyrin, we wondered if hemin may flip B-DNA to Z-DNA as reported for Zn porphyrins (*25*). Figure 1E and F indicate that addition of 5 μM hemin to the growth media (H-TSB) promotes formation of streamers by *S. epidermidis* 1457 grown in TSB with or without 200 mM NaCl (H-TSB-NaCl). The same effect was observed for the clinical isolate *S. epidermidis* AUH4567 (Figure 1G and Figure S1). We also confirmed that streamers only formed when biofilms were grown with mechanical agitation (Figure 1G and H). The involvement of non-canonical DNA structures in streamer formation was further supported by the observation that streamers formed in the presence of NaCl or KCl (Figure 1I), which stabilize GQ and Z-DNA. In contrast, no streamers were seen in the presence of CsCl (Figure 1J), which promotes neither GQ nor Z-DNA, but possibly induces an alternative DNA secondary structure (*42*).

After confirming that 5 μM hemin did not impede planktonic growth (Figure S2), we used TSB with 200 mM NaCl and 5 μM hemin as our standard growth medium to study extracellular DNA structures in *S. epidermidis* biofilm from two clinical isolates, strain 1457 and AUH4567.

### *S. epidermidis* biofilm contains eDNA rich in G-quadruplex and Z-DNA

The next objective was to determine which non-canonical DNA structures were present in the biofilm matrix. We first optimized immunolabelling of biofilms to visualize GQ and total DNA (Figures S3 and S4), and subsequently used full immunolabelling to determine the relative abundance of Z-DNA, GQ, triplex DNA and i-motif compared to B-DNA (Figure 2A-D). A relative quantification of the non-canonical DNA structures was obtained by normalizing the fluorescence from antibodies binding to non-canonical DNA by the fluorescence from antibodies binding to B-DNA within the sample (Figure 2E). Quantification was based on area coverage in 2D confocal laser scanning microscopy (CLSM) images in DAIME (*43*) as well as total fluorescence from fluorometric analysis of entire biofilms in 96-well plates. Among the non-canonical DNA structures, Z-DNA and GQ were the most abundant (Figure 2E). Interestingly, CLSM imaging revealed that these structures were primarily present in web-like strings stretching between clusters of bacterial cells (indicated by arrows in Figure 2A).

**Figure 2.**
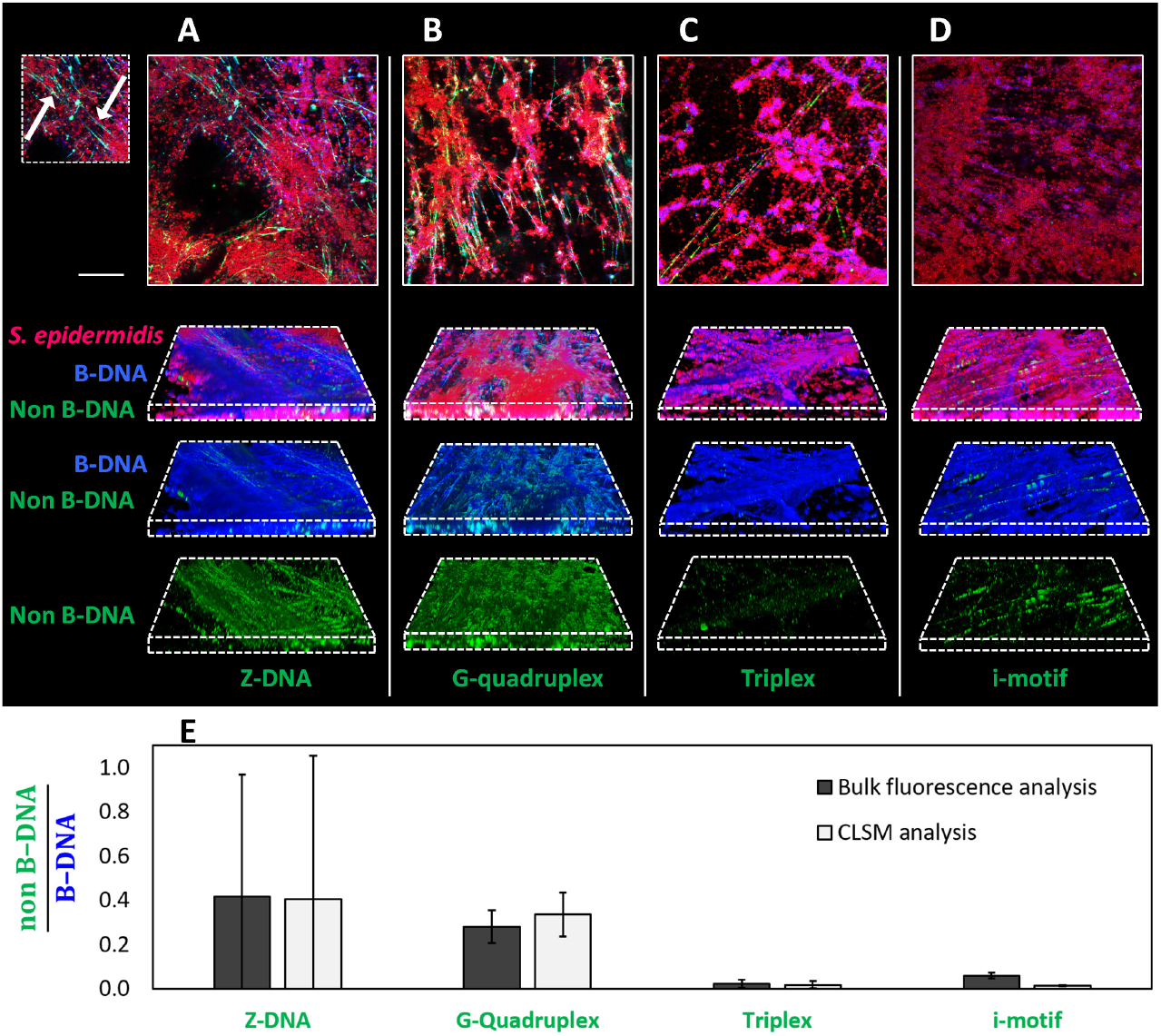
*S. epidermidis* biofilms contain large amounts of Z-DNA and GQ. *S. epidermidis* AUH4567 biofilms grown for 3 days in H-TSB-NaCl with 150 rpm shaking. CLSM images show bacterial cells in red (FM4-64 stain), B-DNA in blue (AB1-AB2 antibody) and non-B-DNA structures in green (antibodies Z22, BG4, Jel466 and iMab for Z-DNA, GQ, triplex DNA and i-motif, respectively). (**A-D**) CLSM 2D and 3D images of the biofilms, using identical acquisition settings. Scale bar = 20 μm. Arrows indicate examples eDNA strings harboring Z-DNA in (A). (**E**) Quantification of non-B-DNA population relative to B-DNA based on area of fluorescence from 2D CLSM images (light grey bar) and bulk fluorescence from Atto488 (immunolabelled non-B-DNA) normalized to bulk fluorescence from CF505S (immunolabelled B-DNA) (dark grey bar). Bars show mean values ± S.D. (n=4).

### Polysaccharides and hemin promote formation of Z-DNA in *S. epidermidis* biofilm

The PNAG polysaccharide is abundant in laboratory-grown *S. epidermidis* biofilm (*8, 44*), and it has been shown to associate with eDNA (*38*), similarly to what has been shown for other cationic polysaccharides in other species (*45*). We wondered if PNAG played a role in the transition of B-DNA to either GQ or Z-DNA. To address this question, we visualized GQ and Z-DNA formed in biofilms lacking PNAG, using *S. epidermidis* 1457 *ΔicaADBC* (Figure 3). The strain produced significantly thinner biofilm compared to the wildtype strain (Figure 1D, F), and CLSM images revealed abundant GQ (Figure 3A, B), while Z-DNA was almost absent (Figure 3C, D) in the polysaccharide-free biofilms. Interaction of DNA with polycations can transition DNA from B- to Z-form (*23*), and the partially deacetylated PNAG may thus prompt this transition due to its polycationic nature. To support this hypothesis, we investigated if chitosan, which is similar in structure to fully deacetylated PNAG, could cause the B to Z-DNA transition *in vitro*, using DNA oligonucleotides (Table S1) whose conformations were monitored by circular dichroism to distinguish between B- and Z-DNA. Indeed, chitosan caused this transition in a dose-dependent manner, resulting in the full transition to Z-DNA at ≥0.025% chitosan in Tris-acetate buffer at pH 5.5 (Figure S5).

**Figure 3.**
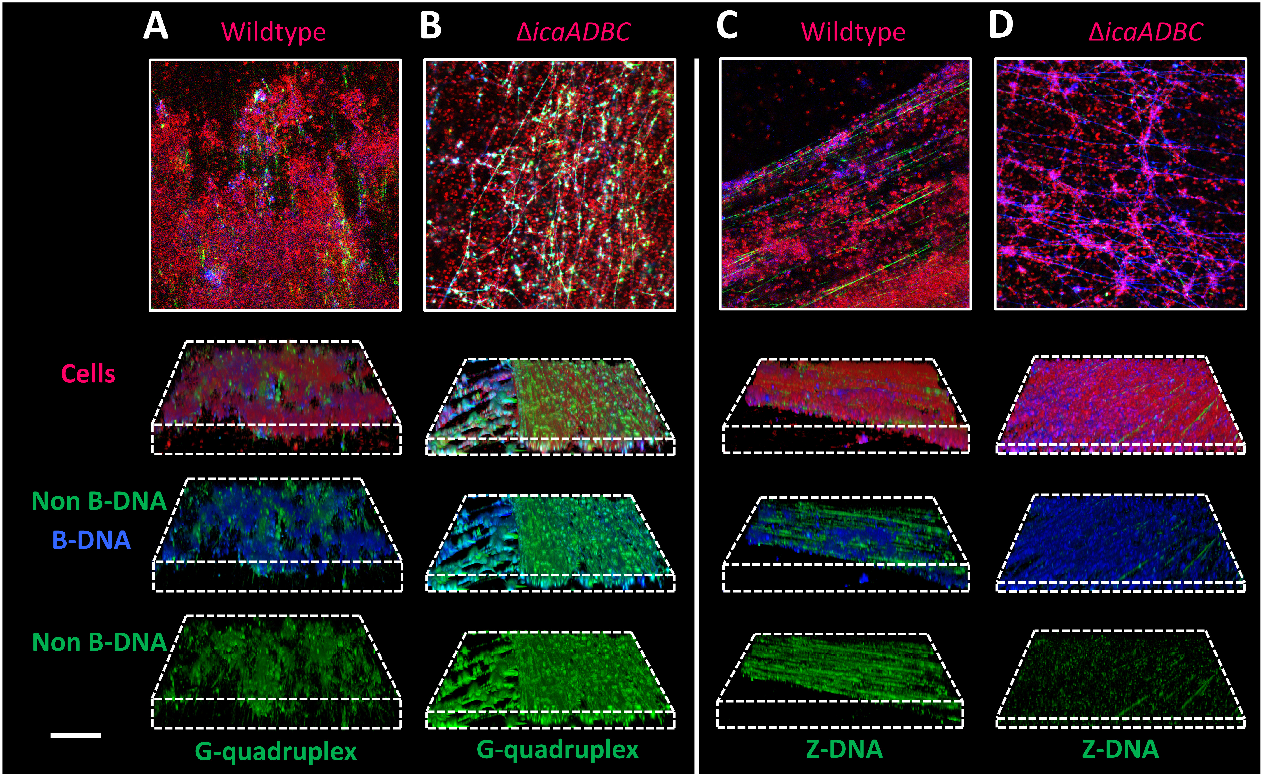
Polysaccharide production stimulates formation of Z-DNA but not GQ. *S. epidermidis* 1457 WT and Δ*icaADBC* mutant strains were grown in H-TSB-NaCl for 3 days (150 rpm shaking). Images show immunolabelling of GQ (A, B) and Z-DNA (C, D) in *S. epidermidis* 1457 wildtype (A, C) and the isogenic polysaccharide-deficient mutant (B, D). 2D images (top panel) and 3D images (bottom panel) show cells in red (FM4-64 stain), B-DNA in blue (AB1-AB2 antibody), GQ (BG4 antibody) or Z-DNA (Z22 antibody) in green. Acquisition settings are identical in all images (same as in the images in Figure 2). Scale bar = 20 μm.

We observed that in addition to the presence of eDNA and polysaccharides, several environmental factors such as cations, hemin, and mechanical stress affected the formation of macroscale streamers in *S. epidermidis* biofilms (Figure 1). We hypothesized that non-canonical DNA structures were involved in streamer formation, and we therefore investigated the effect of hemin, NaCl, and mechanical stress on GQ and Z-DNA formation in biofilms. The results of biofilm immunolabelling suggested that while GQ formation was promoted by NaCl alone (Figure 4A), Z-DNA formation was only prominent in biofilms grown with both NaCl and hemin (Figure 4B). Moreover, the impact of hemin and NaCl was only seen alongside mechanical stress from the 150 rpm rotational shaking during biofilm growth that possibly induced bending of eDNA. Bending of DNA has already been shown to facilitate formation of non-canonical DNA secondary structures under physiological conditions. (*39*)

**Figure 4.**
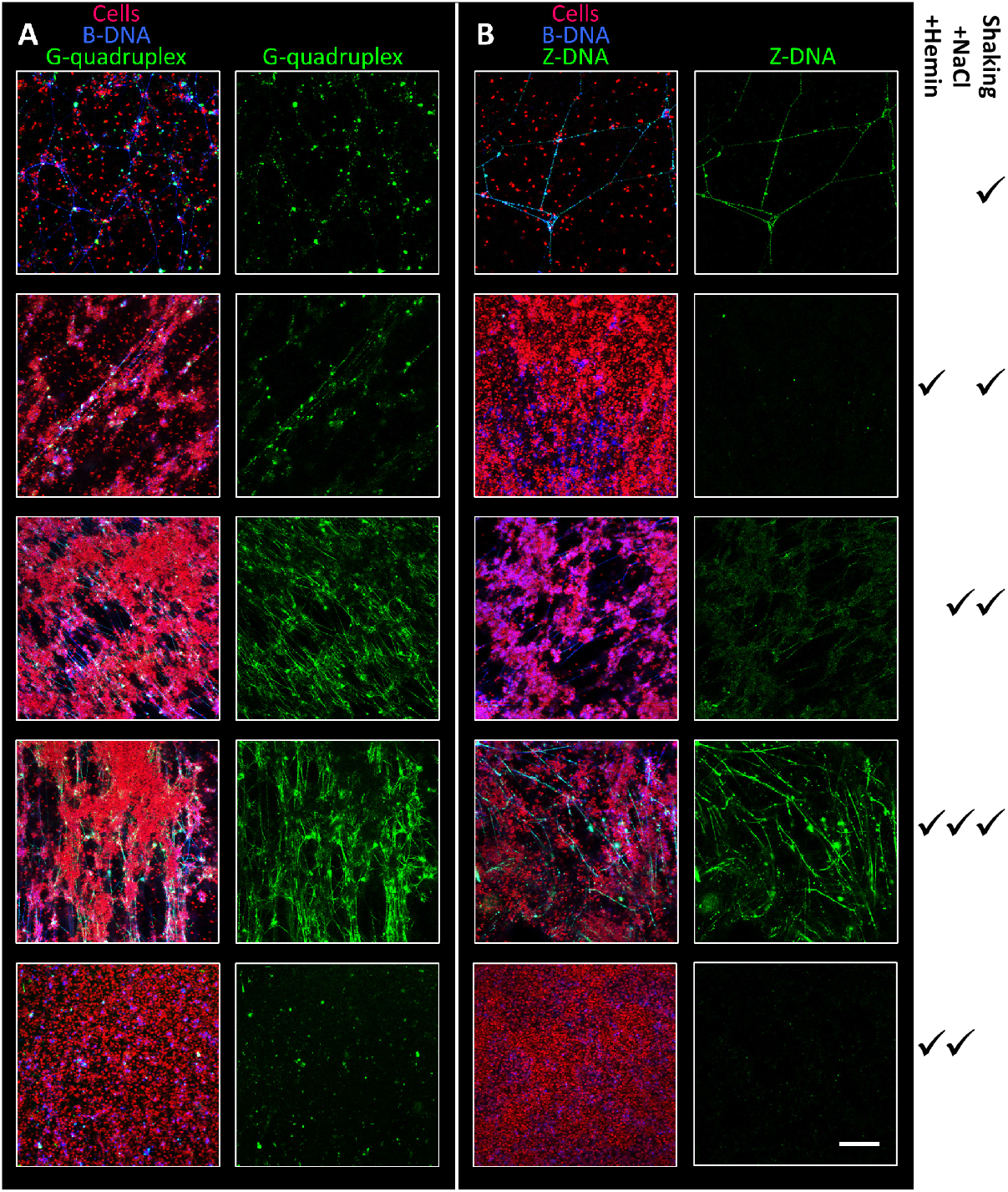
Hemin and NaCl stimulate formation of Z-DNA and NaCl stimulates formation of GQ, but only in combination with mechanical stress from 150 rpm shaking. *S. epidermidis* AUH4567 biofilms were grown for 3 days in TSB with 5 μM hemin or 200 mM NaCl as indicated. Cells are shown in red (FM4-64 stain), B-DNA in blue (AB1-AB2 antibody), GQ (BG4 antibody) or Z-DNA (Z22 antibody) in green. The altering growth conditions are marked by “✓”. Scale bar = 20 μm.

### Hemin binds to G-quadruplex DNA in *S. epidermidis* biofilms and forms a peroxidase-like DNAzyme

DNA and RNA G-quadruplexes bind hemin *in vivo* (*41*) and form a DNAzyme with peroxidase activity similar to horseradish peroxidase (*46*). GQ/hemin DNAzymes have been studied extensively *in vitro* and used in e.g. biosensing, and so far Lat *et al*. provided a first evidence for GQ-hemin-mediated nucleic acid peroxidase activity to work *in vivo* under the right experimental conditions (*46*). We therefore visualized the specific location of DNAzyme activity in *S. epidermidis* AUH4567 biofilms using a reaction similar to tyramide signal amplification (*46*) (Figure 5A) followed by immunolabelling of GQ-DNA and B-DNA to determine if the peroxidase activity co-localized with GQ. As a positive control for this method, we grew biofilms amended 2 μM of the GQ-forming DNA oligo c-myc-4 (Table S1), and the position of peroxidase activity (green) in these biofilms matched the location GQ-DNA visualized by immunolabelling (red) (Figure 5B). The same co-localization of GQ-DNA and peroxidase activity was present in biofilms grown without amendment of additional GQ-DNA (Figure 5C). While c-myc is known to form a parallel GQ-DNA, we expected different types of GQ-DNA to form in the biofilms without c-myc. The different variants of GQ vary in peroxidase activity of the GQ/hemin complex, and this could explain why tyramide deposition was not present at all locations with GQ-DNA. Indeed, the DNAzyme activity is primarily associated with the parallel GQs, and the location of peroxidase activity may thus reflect the location of parallel GQ/hemin complexes (*46*). We noticed that the peroxidase activity as well as BG4-immunolabelling was also found in clustered areas that did not overlap with B-DNA immunolabelling, indicating that these G-quadruplexes may be GQ-RNA (Figure S6).

**Figure 5.**
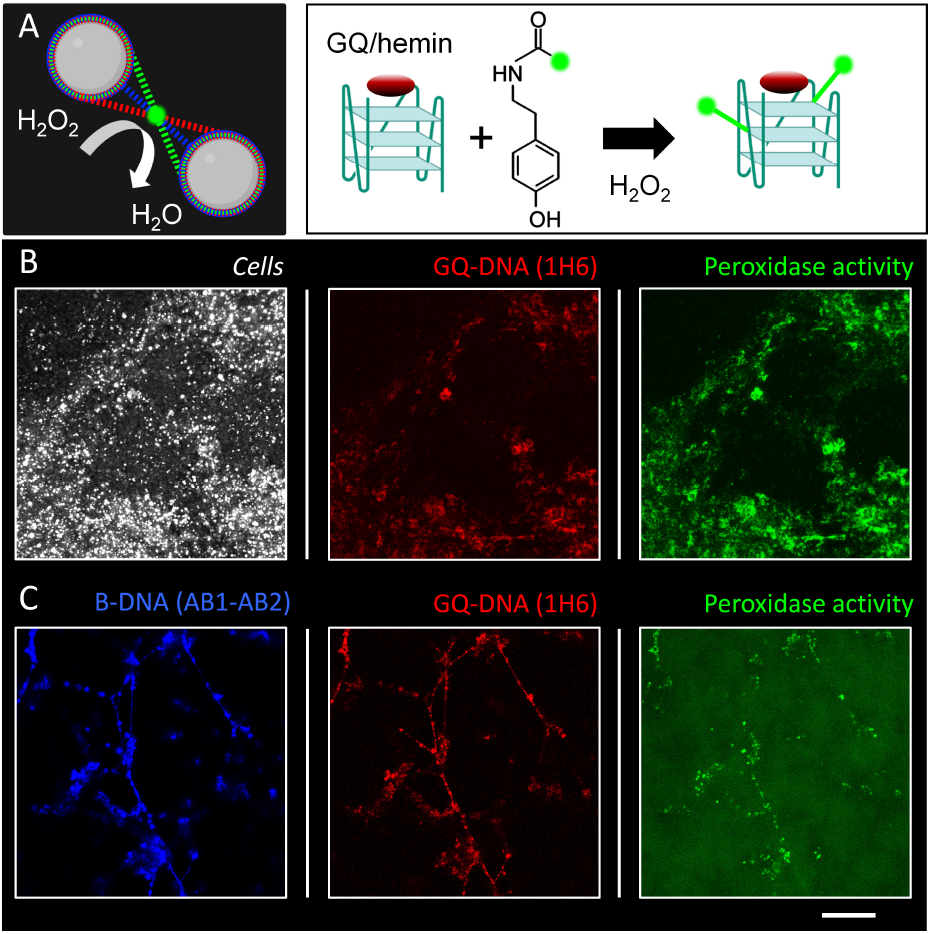
Hemin and GQ enable peroxidase-like DNAzyme activity in biofilms. (A) Schematic presentation and (B) 3D CLSM and (C) 2D CLSM images showing extracellular DNA immunolabelling and tyramide signal amplification (linked to the peroxidase activity) in 3-day *S. epidermidis* AUH 4567 biofilms (H-TSB-NaCl, 150 rpm shaking). (B) Peroxidase (green), GQ-DNA (red) and bacteria (white) labelling in biofilms doped with 2 μM c-myc-5. (C) Peroxidase (green), GQ-DNA (red), and B-DNA (blue) labelling in biofilms without doping. Scale bar 20 μm.

### *In vivo Staphylococcus aureus* biofilm from murine osteomyelitis model contains GQ and Z-DNA

The formation of non-canonical secondary DNA structures in biofilms may be an observation resulting from the artificial conditions imposed on biofilms grown in the lab. We therefore investigated if GQ and Z-DNA were also present in biofilms grown *in vivo*. For this purpose we used *Staphylococcus aureus* biofilms from a murine osteomyelitis model where 18 h biofilms were formed on the implant in TSB *in vitro*, subsequently inserted into the tibia, and left for 7 days before extraction and CLSM imaging. Figure 6 shows the presence of both GQ and Z-DNA in the tissue attached to the implant surface, confirming that non-canonical DNA structures also form *in vivo*. The biofilm did not have any detectable autofluorescence, however, the implant surface had an autofluorescence in green channel (Figure S7).

**Figure 6.**
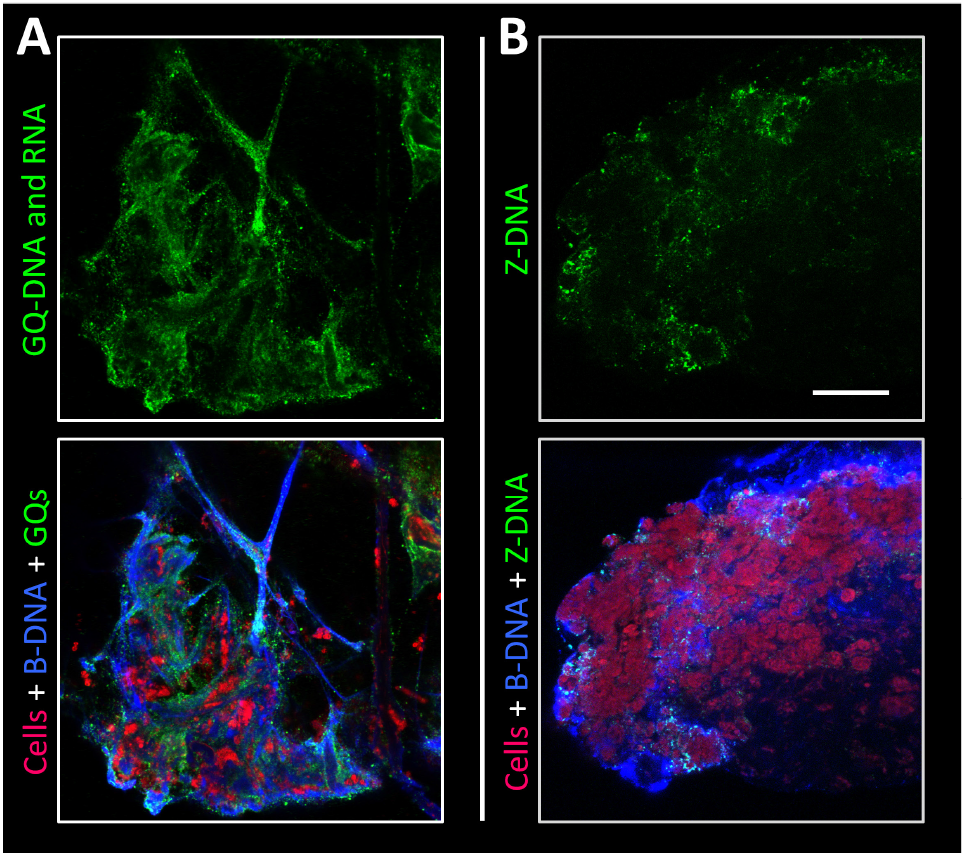
In vivo biofilm from murine osteomyelitis model contains GQ-DNA and Z-DNA. 2D CLSM images of the tissue attached to tibia implants infected with *S. aureus*: (A) GQ and (B) Z-DNA. Cells (bacterial and murine) are shown in red (FM4-64 stain), B-DNA in blue (AB1-AB2 antibody), and GQ (BG4 antibody) or Z-DNA (Z22 antibody) in green. The images are shown as single channel (green) as well as three-channel. Scale bar 20 μm.

### Bacterial but not mammalian nucleases can degrade GQ and Z-DNA

A benefit from forming non-canonical DNA structures in the biofilm matrix could be to protect the biofilm from degradation by host DNases. We therefore investigated if mammalian DNase I or other nucleases could degrade GQ and Z-DNA. To address this question, we developed a fast assay to quantify nuclease activity against GQ-DNA, Z-DNA, ssDNA and B-DNA, and used it to compare double strand-specific DNase I with single strand-specific nucleases of bacteria origin, namely the Micrococcal nuclease originating from *Staphylococcus aureus*, and S1 nuclease originating from *Aspergillus oryzae*. Moreover, micrococcal nuclease is used to degrade total DNA such as chromatin DNA (*47*) and RNA (*48*) in sequencing and mapping assays.

Intra-strand GQ (1 μM) were prepared by folding parallel and hybrid GQ from oligos using c-myc and tel DNA sequences (Table S1), respectively, in 100 mM KCl. Single-stranded DNA oligos “ssB1” and “nonGQ” (G-rich but unable to form intra-strand GQ) and double-stranded DNA oligos “dsB1-B1c” samples were prepared in the same buffer and included as controls. The B-DNA and non-canonical DNA structures were confirmed by circular dichroism (Figure S5). Z-DNA (1 μM) was folded from two complementary DNA sequences in 25 mM Tris buffer, 6.25 mM CaCl_2_ and 1 mM MgSO_4_ (pH 6) with 0.025 % chitosan. We used a random sequence, “Z1”, and a GC-rich sequence, which is assumed to form more stable Z-DNA, “Z2”. For comparison, B-DNA of the same sequences were prepared (B1 and B2) without addition of chitosan. The annealed DNA sequences (in their buffers) were diluted to 200 nM in 25 mM Tris buffer, 6.25 mM CaCl_2_ and 1 mM MgSO_4_ (pH 6) to allow optimal nuclease activity before adding the nuclease or water (blanks) to start the incubation. After incubation with nucleases, fluorescent DNA-binding stains were added to assess the amount of undigested DNA remaining (Figure 7). Due to its superior detection of ssDNA and GQ, we used SYTO60 for DNA quantification in samples where GQ and B-DNA were compared (Figure 7A), while Picogreen was used for DNA quantification in samples where Z- and B-DNA were compared (Figure 7B). The ability of nucleases to degrade the different DNA structures was assessed by normalizing the fluorescence intensity after 2 h enzyme treatment to reference samples that did not receive the enzyme, i.e. a value of 1 reflects no DNA degradation, while 0 reflects full DNA degradation. A single sample had a value >1, which may be caused by increased fluorescence of a DNA-binding dye when DNA-binding proteins bind.

**Figure 7.**
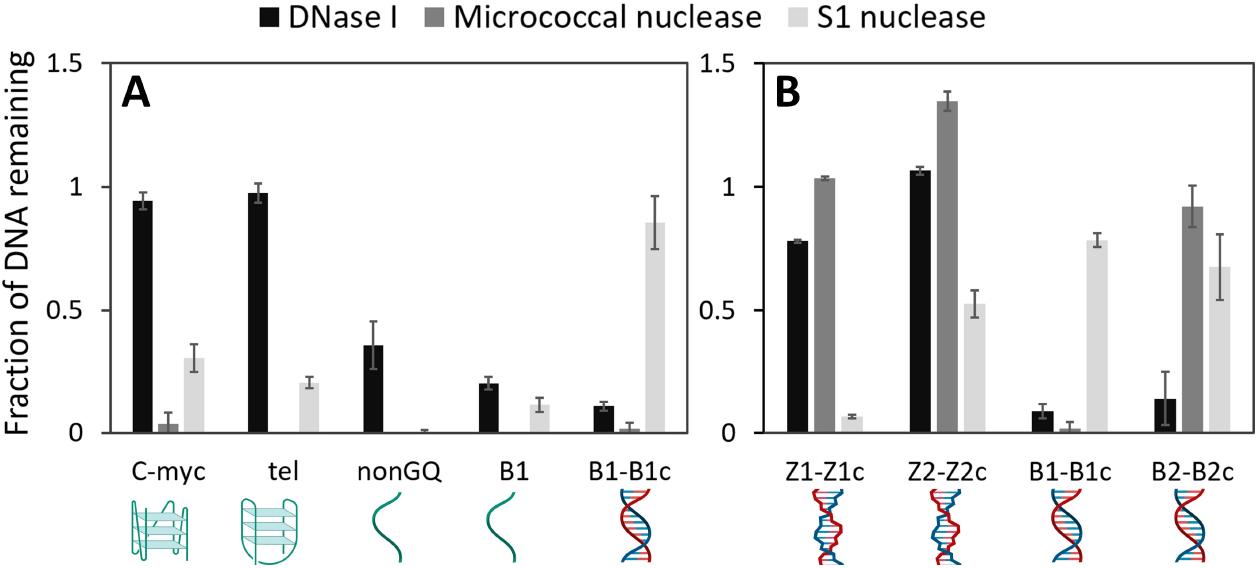
DNase I does not degrade non-canonical DNA structures, while Micrococcal nuclease degrades GQ, and S1 nuclease degrades Z-DNA. Pre-folded oligos (dsDNA, ssDNA, GQ and Z-DNA) were incubated with nucleases for 2 h, and the remaining DNA was quantified and normalized to the untreated control, by quantification of fluorescence from the DNA-binding dye SYTO60 (A) or Picogreen (B).

DNase I degraded dsDNA and ssDNA but had no activity towards GQ and Z-DNA. Micrococcal nuclease fully degraded GQ, ssDNA and dsDNA in the B-DNA form, but had no activity against Z-DNA. S1 nuclease is known to be specific for ssDNA, and as expected, it fully degraded ssDNA but had no activity against dsDNA in the B-DNA form. However, it partially degraded GQ and Z-DNA, particularly the Z-DNA formed by the random “Z1” sequence. Some activity of S1 nuclease toward these structures was expected, as BZ junctions as well as the single-stranded DNA loops connecting GQ in supercoiled plasmids are susceptible to S1 nuclease (*21, 49*). In summary, non-canonical DNA structures are resistant to the action of mammalian DNase I, but nucleases of bacterial origin may be used to tackle these robust DNA structures.

Based on the results of pure DNA *in vitro* assays, we hypothesized that Micrococcal nuclease and S1 nuclease could remove GQ in biofilms. To test this, we treated the *S. epidermidis* biofilms in the same buffer and at the same enzyme concentration as used for DNA oligos, and assessed the eDNA remaining in the biofilm by immunolabelling and CLSM while the overall biofilm removal was assessed by label-free 3D imaging using OCT. CLSM images revealed an overall decrease in eDNA in samples treated with the Micrococcal nuclease (Figure 8A,B), indicating removal of both B-DNA and GQ by this enzyme. In contrast, S1 nuclease had little impact on eDNA in the biofilm (Figure 8A,C), and only showed indication of removing some of the web-like DNA strings. DNase I only removed B-DNA (Figure 8A,D), but quite surprisingly, large areas of aggregated GQ emerged after DNase I treatment, indicating the DNase I resulted in remodeling of the eDNA. The biofilm volume after treatment with Micrococcal nuclease, S1 nuclease and DNase I decreased by 36 %, 6 %, and 15 %, respectively, which was corroborated observations by CLSM (Figure 8E). Collectively, these results suggest that Micrococcal nuclease is more effective than DNase I for implementation in biofilm control.

**Figure 8.**
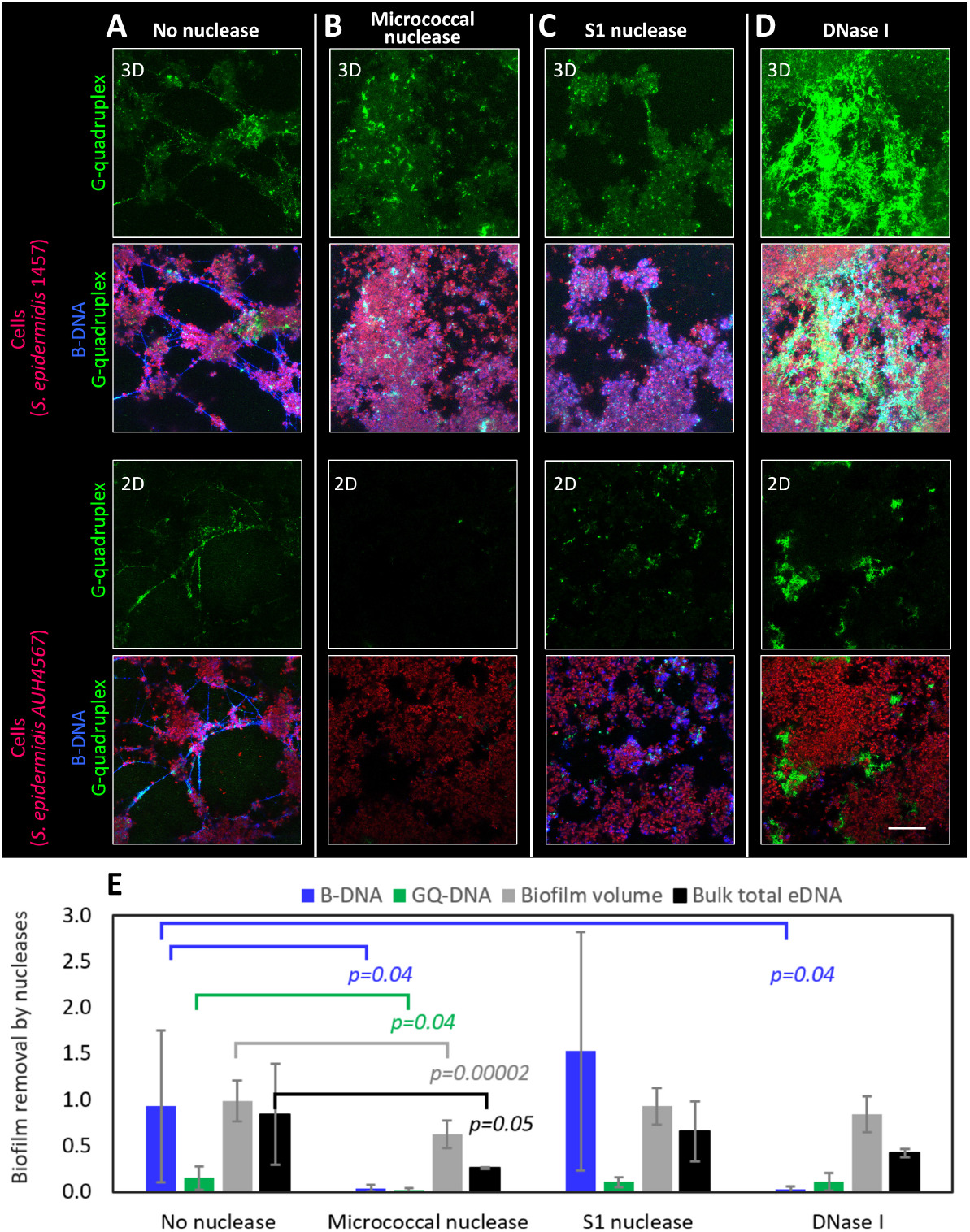
CLSM and OCT imaging of *S. epidermidis* biofilms treated by different nucleases show that Micrococcal nuclease is more effective than S1 and DNase I. *S. epidermidis* AUH 4567 and 1457 biofilms were grown in H-TSB-NaCl for 3 days and treated with (A) no enzyme, (B) Micrococcal nuclease, (C) S1 nuclease, and (D) DNase I in 25 mM Tris-Acetate buffer (pH 6) amended with 6.25 mM Ca^2+^ and 1 mM Mg^2+^. Cells are shown in red (FM4-64 stain), B-DNA in blue (AB1-AB2 antibody), GQ (BG4 antibody) or Z-DNA (Z22 antibody) in green. Acquisition settings are identical in all images. Scale bar = 20 μm. (E) Quantification of eDNA- and biofilm removal by the nucleases. Analysis of eDNA is based on 2D CLSM analysis of B-DNA (blue bar) or GQ (green bar) normalized to the FM 4-64 signal from cells in the CLSM images. Values are mean ± S.D (n = 6). Analysis of biofilm volume (mm^3^, grey bar) is based on 3D OCT imaging. Values are mean ± S.D (n = 15). Analysis of total eDNA in the biofilms (black bar) based on the bulk TOTO-1 fluorescence measured in the plate reader (n = 4) and normalized to the “No nuclease” sample initial fluorescence.

## DISCUSSION

Most biofilms contain eDNA comprised of genomic DNA originating from lysed bacteria or from immune cells that respond to an infection. Outside the controlled environment of a cell, eDNA exists in a variable chemical environment and will interact with other polymeric substances in the biofilm matrix. These interactions and local conditions can cause DNA to adapt non-canonical secondary structures, which may provide new functions and properties of the biofilm matrix, which were previously overlooked. In this study, we show large amounts of GQ and Z-DNA in the extracellular matrix of *S. epidermidis* biofilms grown in laboratory medium amended with hemin and NaCl (Figure 9). Although NaCl concentration used in our study was above physiological levels, we found the same structures in biofilms from implant-associated staphylococcal infections. Furthermore, biofilms experience much higher concentrations of NaCl and KCl e.g. on the skin, or in the marine environment. It is thus likely that biofilms in many different environments also contain non-canonical DNA structures.

**Figure 9.**
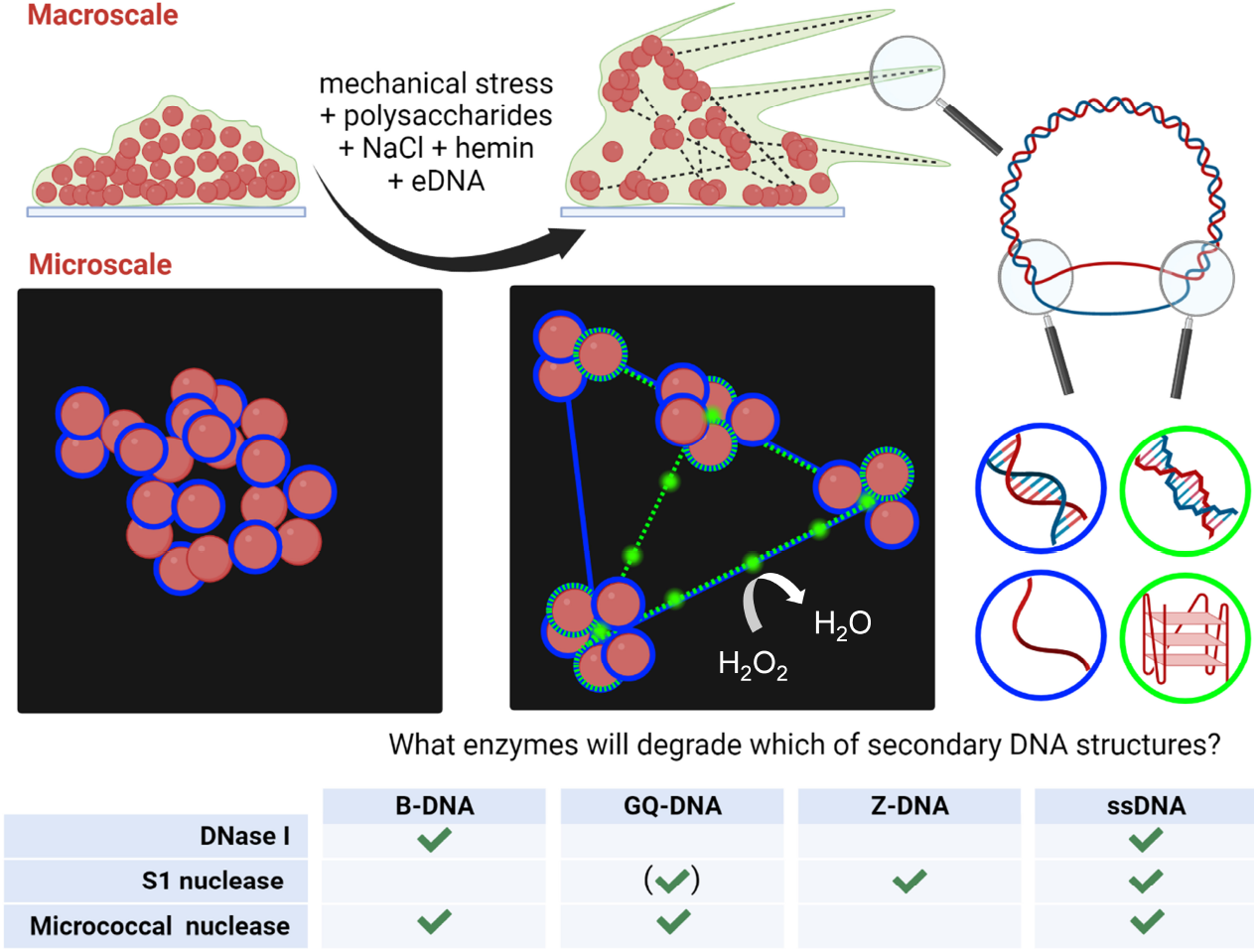
Factors inducing eDNA secondary structures and their roles in *S. epidermidis biofilm*. Environmental and bacteria-driven factors shaping macroscale structure of the biofilm into streamers contribute to formation of non-canonical DNA secondary structures detected at the microscale by immunolabelling and CLSM. GQ-DNA and possibly GQ-RNA scavenge hemin and gain catalytic peroxidase activity. The table summarizes the result of assays quantifying degradation of synthetic GQ- and Z-DNA oligos by different nucleases. The Micrococcal nuclease degraded both GQ-DNA as well as B-DNA and was the only one nuclease that re-dispersed the biofilm.

### Formation of Z-DNA in biofilms

While GQ was formed merely by elevating the NaCl concentration, formation of Z-DNA also required two other components: the polysaccharide PNAG and hemin (Figure 3 and 4). The role of hemin in the B-to-Z-DNA transition is intriguing. We speculate that hemin can react with the eDNA, turning guanines into 8-oxoguanines which favor Z-DNA (*28*), or that hemin forms a coordination bond with the N7 atoms of guanines as it has been proposed for the B-to-Z transition mediated by Zn porphyrins (*25*). While there is no evidence of hemin forming coordination bond with N7 atom of guanines, a similar coordination bond has been reported between heme and histidine- and tyrosine-based protein motifs (*50*), supporting that such coordination is a plausible mechanism for hemin’s role in Z-DNA formation. *In vivo*, bacteria have access to hemin in plasma (2-5 μM (*51*)) or on inflamed mucosal surfaces, as exemplified by the requirement for hemin (≈ 7.5 μM) by some biofilm-forming pathogens (*52*). We confirmed that both GQ and Z-DNA were present *in vivo* in implant-associated *S. aureus* biofilms from a murine model (Figure 6), and a previous study of Z-DNA also found large amounts of Z-DNA in *in vivo* biofilms from lung-or middle ear infections as well as in *in vitro* biofilms that were grown with hemin (*17*).

The involvement of extracellular polysaccharides in Z-DNA formation is most likely linked to their polycationic nature. PNAG in *S. epidermidis* biofilms is approximately 15-20% deacetylated (*53*) resulting in a net positive charge at neutral pH. Polycations can promote B to Z-DNA transition (*23*), which we also confirmed in a model system using chitosan as the polycationic agent (Figure S5). Bacteria produce several different polycationic matrix components. These include e.g. other positively charged polysaccharides, such as Pel in *Pseudomonas aeruginosa* which also co-localizes with eDNA (*45*), or polycationic peptides, such as the α-type phenol-soluble modulins which also complexes with eDNA in biofilms (*54*). We also note a large variation in Z-DNA abundance between different samples (Figure 2), and Z-DNA was primarily present in web-like network of DNA strings while it was largely absent from the eDNA associated with the surface of bacterial cells (Figure 3 and 4). One of the factors driving formation of non-canonical DNA structures can be unzipping of B-DNA due to bending (*39*), and the location of both GQ and Z-DNA in web-like strings suggests that a mechanical force could affect their formation in biofilms. No such strings were observed in biofilms grown without mechanical stress from 150 rpm rotational shaking (Figure 1 and Figure 4).

In addition to the environmental factors and biological components discussed here, Buzzo *et al*. also confirmed the involvement of DNA-binding proteins of the DNABII family, which are universally present in bacteria and are believed to promote Z-DNA formation by bending DNA and by stabilizing Holliday junctions between DNA strands in the biofilm matrix (*17*). Collectively, the B- to Z-DNA transition of eDNA in biofilms is a result of specific environmental conditions, and the presence of hemin and bacteria-derived matrix components, which most likely differ between bacterial strains.

### Formation of GQ in biofilms

G-quadruplex formation in biofilms could involve both extracellular DNA and RNA and can either be intra-stranded (one strand) and inter-stranded (2 or 4 strands) with conformations known as parallel, hybrid or anti-parallel. The specific form depends on the variety of environmental conditions, e.g. supercoiling (*29*), crowding (*32*), type and concentration of divalent (*33*) and monovalent (*34*) cations and the GC content of nucleic acid itself. The only previous study to report G-quadruplexes in biofilms identified GQ-DNA and GQ-RNA in extracellular nucleic acids extracted from *P. aeruginosa* biofilms (*18, 19*). Our study is therefore the first to visualize these structures directly in the biofilm. While the 1H6 antibody detects primarily GQ-DNA (*18, 55*), the BG4 antibody used in our study detects both GQ-DNA and GQ-RNA (*56*). The binding of both 1H6 and BG4 antibodies to the total eDNA revealed 1H6 bound in discrete spots while the signal from BG4 was brighter and more continuous along the web-like DNA strings (Figure S3). Hence, we cannot exclude the contribution of GQ-RNA. However, the antibody signal overlapped with the signal from B-DNA binding antibody, and we therefore believe that the signal is primarily from GQ-DNA.

### Resistance of non-canonical DNA to degradation by nucleases

A potential benefit from having non-canonical DNA structures in the biofilm matrix is their resilience towards DNase I, and this property might explain the numerous reports of DNase I failing to degrade mature biofilms (*14, 17*). Furthermore, eDNA in biofilms grown at high salinity have been reported to be resistant to degradation by DNase I (*16*). For the first time, we address the effect of different nucleases on B-DNA and GQ in biofilms after showing different activities to B-DNA, Z-DNA and GQ-DNA by three nucleases (Figure 7). While our results confirmed that DNase I is specific to B-DNA, we also discovered that DNase I treatment resulted in formation of large GQ aggregates in the biofilm, which were not observed prior to DNase I treatment (Figure 8). The most effective DNase was Micrococcal nuclease, which was the only DNase that reduced the biofilm volume (Figure 8).

The efficacy of bacterial nucleases toward GQ-DNA and Z-DNA (Figures 7 and 8) may reflect evolution of nucleases that enable biofilm dispersal while host nucleases have no impact on the biofilm. DNase I prefers a double-stranded helix as a binding site though it can degrade both double- and single-stranded DNA (as it cleaves one of the complementary strands at a time). However, DNase I is inefficient at cleaving loops (*57*). In contrast, both S1 nuclease and Micrococcal nuclease preferentially cleave single-stranded regions, in the loop or at the ends of a hairpin (*57*). Indeed, the conformational junctions between Z-DNA and B-DNA in supercoiled plasmids are known to be susceptible to the ssDNA-specific S1 nuclease originating from *Aspergillus* (*21, 49*), and synthetic GQ-DNA has been reported to be degraded by Micrococcal nuclease originating from *Staphylococcus aureus* (*58*). We note that being primarily ssDNA-specific nuclease, the Micrococcal nuclease did not degrade dsB2-B2c (100 % GC).

### DNAzyme activity in the biofilm matrix

Another benefit of GQ may be linked to their ability to form a DNAzyme. For the first time, we visualized GQ/hemin peroxidase activity in a biofilm using tyramide signal amplification (Figure 5). Peroxidases are usually considered hazardous to bacteria and are natural antimicrobial components of milk, saliva, and released by immune cells, and are used as antimicrobial components in healthcare and food products (*59*). So how might biofilms benefit from having DNAzymes immobilized throughout the extracellular matrix? One benefit might be protection against certain antimicrobials. Peroxidases can break thioether bonds (*60*) in common antimicrobial peptides, such as nisin, and thereby contribute to the biofilm’s resilience.

### Conclusion

In conclusion, we revealed that *S. epidermidis* produces a thick biofilm matrix that contains web-like structures of eDNA with two non-canonical secondary structures: G-quadruplexes and Z-DNA. Formation of these structures is promoted in the presence of salts (Na^+^ and K^+^) that stabilize the structures, and also in the presence of the metalloporphyrin hemin. Furthermore, mechanical stress that pulls at the eDNA is also involved in their formation.

The potential advantages for forming non-canonical DNA structures in the biofilm matrix includes increased mechanical strength and protection against degradation by certain nucleases. Furthermore, GQ/hemin complexes provide sites of peroxidase activity in the extracellular matrix and represent a novel emergent property of the biofilm matrix with potentially protective properties against antimicrobial peptides.

## METHODS

### Chemicals and buffers

Stock solutions of 1 M Tris hydroxymethyl aminomethane (Tris pH 7.5, Merck), 2 M NaCl (Merck), 2 M KCl (Merck), 2 M MgSO_4_ (Merck), 670 mM CaCl_2_ (Merck) were prepared in milliQ water (18.2 Ω). The pH was adjusted with 33% acetic acid (Merck). A 2 % chitosan solution was obtained by dissolving chitosan (high purity, 740063 Merck) in milliQ water with 0.13 % (vol/vol) acetic acid followed by repeated cycles of heating to 50°C for 5 minutes and vortexing for 30 s. A 25 mM hemin solution was obtained by dissolving hemin (51280-1G, Merck) in 200 mM Tris with 100 mM NaOH (Merck) and 30 % DMSO (pH 11). Hemin was handled and stored away from light. A 1 × GQ-buffer consisted of 10 mM Tris with 100 mM KCl (pH 7.5) and was used for annealing GQ from DNA oligos. A 1 × B-buffer consisted of 25 mM Tris with 6.25 mM CaCl_2_ and 1 mM MgSO_4_ (pH 5.5) and was used for annealing B-DNA and for all DNase treatments. 1 × Z-buffer consisted of 25 mM Tris with 6.25 mM CaCl_2_ and 1 mM MgSO_4_ and 0.025 % chitosan (pH 5.5) and was used for annealing Z-DNA. All solutions were sterile filtered using 0.2 μm filters (83.1826.001, Fisher Scientific).

### Fluorescent stains

Picogreen Quant-iT™, TOTO-1, TOTO-3, SYTO60 and FM 4 64 (Thermo Fisher Scientific) were aliquoted to 100 × working concentration in milliQ water. DNA-binding stains were stored at -20 °C and FM 4-64 was stored at 4 °C.

### Synthetic DNA substrates

DNA oligos (Table S1) were purchased from Integrated DNA Technologies (IDT) as stocks of 100 μM in 1 mM TE buffer (pH 7.5), purified using standard desalting. Solutions of DNA in either 1 × GQ-buffer (10 mM Tris, 100 mM KCl, pH 7.5) or in 1 × Z-buffer (0.025 % chitosan, 25 mM Tris, 6.25 mM CaCl_2_, 1 mM MgSO_4,_ pH 5.5) or 1 × B-buffer (25 mM Tris, 6.25 mM CaCl2, 1 mM MgSO4, pH 5.5) were annealed to obtain pure GQ-proxy, Z-proxy or B-proxy model, respectively. For circular dichroism (CD), 5 μM DNA solutions were annealed to obtain better quality of the signal. For the enzymatic assays, 1 μM DNA solutiuons were annealed.

### Characterization of DNA substrates by Circular Dichroism

The structure of Z-DNA was verified with the use of CD signal measured on J-810 Spectropolarimeter with wavelength range at 220-320 nm, data pitch at 0.5 nm, scanning speed at 50 nm/min, response at 4 sec and bandwidth at 2 nm. The measurements were done by loading 60 μL of the sample to a cuvette with 0.3 cm optical path. Samples of 5 μM DNA were prepared in 1 × B-buffer doped with 0-0.1 % chitosan and annealed as described previously. The signal measured was CD (mdeg = 0.001 deg), which is related to absorbance by a factor of 32.98. Taking this into account, we converted the CD signal to Δε(M^(−1) cm^(−1)) with use of Lambert-Beer law:

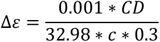

The DNA concentrations were calculated in moles of base pairs per liter by multiplying 5E-06 M concentration by the number of base pairs in the two DNA sequences analyzed.

### Anti-DNA antibodies

The 1 mg/mL antibodies Atto488-BG4 (goat monoclonal IgG lambda), FluoProbes647®-1H6 (goat monoclonal IgG kappa), and Atto488-Z22 (rabbit monoclonal IgG kappa) in phosphate buffer saline (PBS) with 0.02 % proclin were purchased from the Absolute Antibodies as fluorescently tagged antibodies (subsequently stored in the fridge with light protection). The 1 mg/mL antibody AB1 (mouse monoclonal 3519) in PBS was purchased from Abcam (aliquoted and stored at -20°C). The 2 mg/mL anti-mouse secondary antibody CF405S-AB2 (goat polyclonal IgG H+L) and untagged 1H6 (mouse monoclonal IgG2bκ) in PBS with 0.05 % sodium azide and 50 % glycerol were purchased from Merck (stored at -20°C with light protection). We used sterile-filtered 3 % bovine serum albumin (BSA heat shock fraction, pH 7, > 98 %, Merck) in 1 × Pierce PBS buffer (Thermo Fisher Scientific) to prevent unspecific immunolabelling.

### Media, strains, cultures and plates for in vitro biofilms assays

Autoclaved BHI (Merck, 53286) 37 g/L was used for agar plates (15 g/L agar in BHI) as well as overnight cultures (16-20 h at 37°C with 180 rpm shaking). Autoclaved TSB (Merck, T8907) 30 g/L supplemented with monovalent salts ± hemin was used to obtain biofilms for studying secondary structures of eDNA and their role in biofilm formation. We used the clinical isolates *S. epidermidis* 1457 (WT, Δa*tlE*, and Δ*icaADBC*) (kindly donated by Prof. Holger Rohde, Universitatsklinikum Hamburg-Eppendorf, Hamburg, Germany) and *S. epidermidis* AUH4567 WT (*61*) strains streaked onto tryptic soy agar (TSA) and kept in the fridge for up to 1 month. *In vitro* biofilms were inoculated from overnight cultures diluted 20 times in TSB into 96-well plates, incubated at 37°C at 150 rpm for 6 hours, and subsequently exchanging the media every 20-24 hours. The media used was: (i) TSB, (ii) TSB with 5 μM hemin (H-TSB), (ii) TSB with 200 mM NaCl (TSB-NaCl), (iv) TSB with 200 mM NaCl + 5 μM hemin (H-TSB-NaCl), (v) TSB with 200 mM KCl (TSB-KCl), and (vi) TSB with 200 mM KCl + 5 μM hemin (H-TSB-KCl), (vii) TSB with 200 mM CsCl (TSB-CsCl), and (viii) TSB with 200 mM CsCl + 5 μM hemin (H-TSB-CsCl). Ibidi transparent flat bottom microplate (1.5 polymer coverslip, ibiTreat tissue culture treated, sterile, square well, Ibidi #89626) was used to grow and treat biofilms for confocal laser scanning microscopy (CLSM). Nunc black flat bottom microplate (Nunclon Delta-Treated, sterile, round well, Thermofisher Scientific 137101) was used as substrate to grow and treat biofilms for optical coherence tomography (OCT).

### Murine osteomyelitis in vivo biofilm

This study was approved by the Danish Animal Experiments Inspectorate under permission 2022-15-0201-01133 and was carried out under the supervision of the veterinarians at the Institute of Biomedicine, Aarhus University.

*Staphylococcus aureus* SAU060112, was used for inoculation of the steel implants. An overnight culture was prepared in tryptic soy broth (TSB) media from a single colony of *S. aureus* and incubated for 18 h at 37°C, 180 rpm. The overnight culture was diluted to OD600=0.1 (5 × 10^6^ CFU/mL) in fresh TSB and 5 mL were added to falcon tubes with stainless steel implants (Ento Sphinx, Pardubice IV, Czech Republic) and incubated for 18 h at 37°C.

An osteomyelitis infection was established in mice as previously described (*62*). A total of five 8-10 week old C57bl/6j mice (Janvier Labs, Le Genest-Saint-Isle, France) were included in the study. All animals (n=5) were housed in a IVC (ventilated) cage, at the animal facilities at the Animal Facilities at the Department of Biomedicine, Arhus University at standard room temperature, 12 h day/night cycle with free access water and food. After an acclimatization period of one week, the mice were sedated with isoflurane inhalation anaesthesia (5% induction and 2% cont.) followed by injection of buprenorphine (0.05 mg/kg s.c.) for analgesia. The left hind leg was shaved and the steel implants were implanted through the cortical part of the tibial bone. The implants were bent in a U-shape and cut as close to the skin as possible, and the skin was then manipulated to cover the implant. Mice were returned to their cages and analgesics (buprenorphine, 0.7 mg/kg) were administered to the drinking water the first four days of infection. Following seven days of infection, animals were euthanized under isoflurane anaesthesia by cervical dislocation, and the implants were removed for CLSM imaging.

### Labelling eDNA using anti-DNA antibodies and fluorescent stains

To reduce unspecific binding, biofilms were treated in 3 % BSA in 1 × PBS. After a short blocking step, first, *in vitro* biofilms in separate microwells were incubated with 60 μL solutions of BG4, Z22, Jel466 or iMab antibody (1 : 100) in the 3 % BSA for 75 min. Second, we added 60 μL solution of mouse AB1 antibody (1:100) in the 3 % BSA and continued incubation of the biofilms with the two antibodies for another 75 min. Third, after washing the biofilms with 120 μL of the 3 % BSA, the biofilms were incubated with 60 μL solution of anti-mouse AB2 antibody (1:150) in the 3 % BSA. The biofilms were washed in the 3 % BSA and, finally, stained in 10 mg/L solution of FM 4 64 in 100 mM NaCl. The Atto488-conjugated BG4 and Z22 antibodies were excited by 488 nm laser power 4.5 %, and emitted light was collected at < 600 nm. The CF405S-conjugated AB2 antibody was excited by 405 nm laser power 4.5 % and emitted light was collected at < 550 nm. The FM 4-64 stain was excited at 488 nm laser power 2%, and emitted light was collected at > 600 nm. The detected fluorescence was assigned three colors: blue for CF405S, green for Atto488, and red for FM 4-64.

The samples were visualized as three-channel 2D images as well as maximum intensity projections of Z-stacks (3D images). Cells, B-DNA, and non-canonical DNA were quantified as area coverage in 2D images using the software Daime (*43*).

### GQ/hemin peroxidase activity

For probing the GQ-DNAzyme peroxidase-like activity, we used the tyramide signal amplification to deposit fluorophore-conjugated tyramide at the site of activity. Three-day biofilms grown in H-TSB-NaCl were washed thoroughly in 100 mM NaCl and subsequently incubated with fluorescent Alexa488-tyramide reagent (B40953, Thermo Fisher Scientific) diluted 1 : 100 in 1 × PBS buffer (pH 7.5) with 0.1 % hydrogen peroxide (stock concentration 30 %, Merck) and 2 mM ATP (Thermo Fisher Scientific) at room temperature for 1 h. Subsequently, bacterial cells, B-DNA, and GQ were visualized by immunolabelling and FM4-64 staining as described above.

### Optical Coherence Tomography

We employed OCT to visualize the 3-day *in vitro* biofilms obtained in different media (i) – (viii) (see above) without and with nuclease treatment to provide end-point macroscopic image of the web-like matrix, to identify which media constituents affect the formation of these structures, and whether nucleases can degrade it. The Nunc microwells with the biofilms as well as the spaces between the wells were filled with 100 mM NaCl to the top, and a giant glass coverslip (Caspilor) was placed above the liquid gently to avoid bubble formation. An excess of liquid was removed by a tissue to avoid tilting of the glass. Biofilms were scanned as 500 slices and analyzed using a SD-OCT Ganymede™ 620C1 (Thorlabs GmbH, Lübeck, Germany), with a LSM03 objective lens. Images were acquired with a frequency of 100 kHz and an A-scan averaging of 3. The center of the well (5 mm × 5 mm × 1.49 mm) was imaged with a voxel size of 12 μm × 12 μm × 1.45 μm in the x, y and z dimensions respectively. The biofilm thickness was calculated by a custom written Python script from the two-dimensional B-scans (Supplementary Script). The images were passed through a median filter with a disk-shaped structuring element of radius 2 before image segmentation. The pixels belonging to the biofilm were identified by removing the low-intensity background and high-intensity signal from the plastic substrate by thresholding. The high-intensity pixels of the substrate were used to identify the bottom of the biofilm. The thickness was calculated as the average distance between the top of the biofilm to the plastic substrate, thus including void space within the biofilm.

### Screening nucleases against DNA secondary structures

Three commercial nucleases, i.e. DNase I (4716728001, Merck), S1 (N5661-50KU, Merck), and Micrococcal nuclease (EN0181, Thermofisher) were screened using the pre-fold DNA substrates (Table S1) as well as universal custom 1X B-buffer. The protein concentration was optimized to match activity of the three nucleases: 500 u/mL DNase I, 25 u/mL S1 nuclease and 15 u/mL micrococcal nuclease.

We first tested DNA degradation using 200 nM pure DNA substrates in the in 1 × B-buffer, pH 6, doped with either nuclease or water. The mixtures (180 μL) were assembled directly in Nunc 96-well microwell plate (round, black bottom, untreated, Thermofisher Scientific), mixed, sealed (to avoid evaporation) and incubated at 37°C for 2 hours. At the end, 20 μL of the 10 × Picogreen solution or 20 μL of 10 μM SYTO60 was added to each well containing Z-DNA or GQ-DNA, respectively. For comparison, the stains were also added to ds B-DNA and ss DNA treated the same way (with or without nuclease). The DNA degradation efficiency was then evaluated by quantification of Picogreen fluorescence (ex. 480 nm, em. 520 nm) for ds B-DNA and Z-DNA, or SYTO60 fluorescence (ex. 630 nm, em. 670 nm) for GQ-DNA, ss DNA, and ds B-DNA in a CLARIOstar plate reader (BMG Labtech). Due to variations in fluorescence between samples, we normalized fluorescence against the untreated control samples, i.e. a value of 1 indicates no degradation and a value of 0 indicates full degradation.

The nucleases at the same concentrations (stated above) were used for evaluating nucelase activity against eDNA in biofilms. To investigate eDNA degradation by these nucleases, they were applied directly to the biofilm (grown for 3 days in H-TSB-NaCl at 150 rpm) in the same universal buffer (25 mM Tris buffer with 6.25 mM Ca^2+^ and 1 m Mg^2+^, pH6) and tested at the end-point using CLSM and OCT (see the methods description above). After 3-hour enzymatic reactions, the biofilms were labelled using the standard steps of immunolabelling.

## Supporting information

Electronic Supplementary Information

## ACKNOWLEDGEMENTS

We thank the Villum Foundation (Grant no. 00028321) for funding the major part of this work and Villum Foundation (Grant no. 00050284) for support to complete the project. We also thank Dr. Thomas Seviour for discussions about extracellular RNA in biofilms, and Dr. Holger Rohde for providing the bacterial strains *S. epidermidis* 1457 and isogenic mutants. At last, we thank Dr. Ebbe Andersen for providing us with an access to the fluorescence measurements at the CLARIOstar plate reader.

## Notes

### Competing Interest Statement

The authors have declared no competing interest.

