## Supplementary material for "Extracellular G-quadruplex and Z-DNA protect biofilms from DNase I and forms a DNAzyme with peroxidase activity": Electronic Supplementary Information

### *S. epidermidis* biofilm form a web-like extracellular matrix in the presence of hemin and NaCl

Figure S1 illustrates the web-like biofilm matrix obtained in 3 biological replicates of *S. epidermidis* AUH4567 wildtype in TSB-NaCl vs. H-TSB-NaCl (3-day, 150 rpm shaking) media. Thus, addition of 5 µM hemin to the growth media promoted formation of streamers by both *S. epidermidis* 1457 as well as AUH4567 grown in TSB with 200 mM NaCl (H-TSB-NaCl).

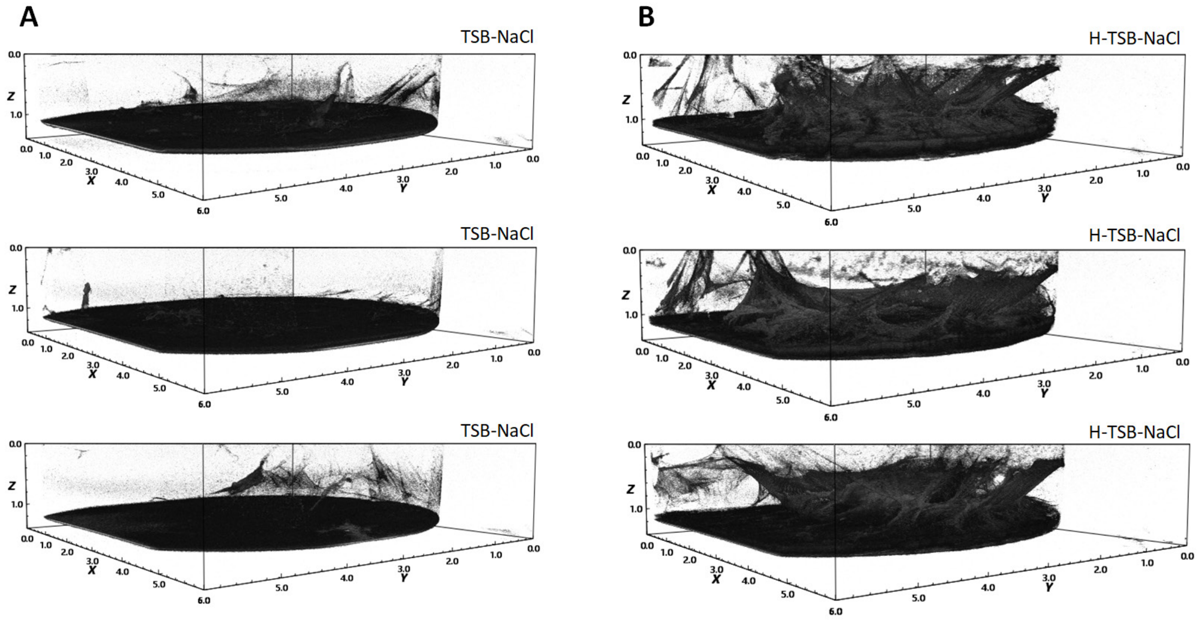


**Figure S1. Hemin induced streamers in 3 days old *S. epidermidis* AUH4567 biofilms**. Three replicates are shown for (A) the WT in TSB-NaCl and (B) the WT in H-TSB-NaCl. Dimensions are 6 x 6 x 1 mm. See Figure 1 in the main manuscript for *S. epidermidis* 1457 biofilms.

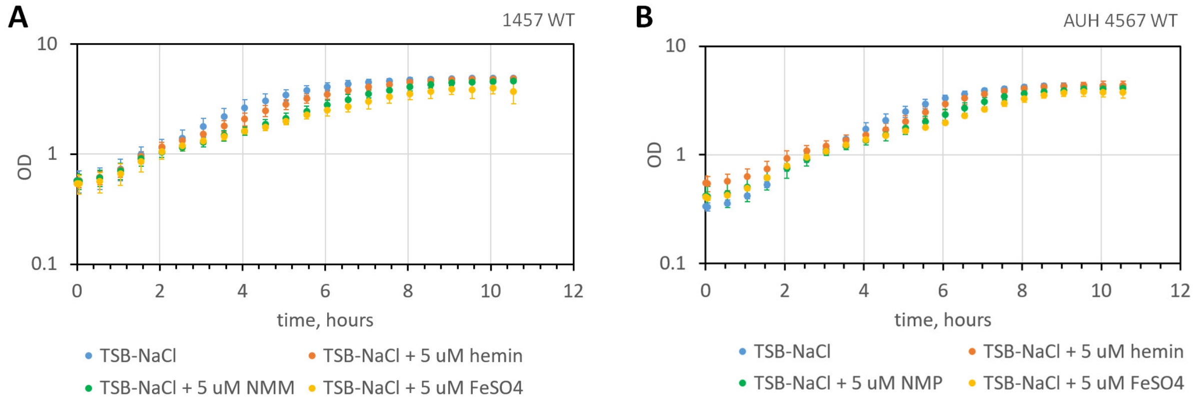


**Figure S2. *S. epidermidis* planktonic cultures grow to OD ≈ 5 in TSB-NaCl as well as H-TSB-NaCl.** Growth curves of *S. epidermidis* 1457 (A) and AUH 4567 (B) in TSB media amended with 5 µM hemin, 5 µM NMP or 5 µM $FeSO_4$ at 37°C (aerobic with no shaking). Error bars show standard deviations of the mean (N = 6).

We investigated whether hemin had an impact onto planktonic growth of *S. epidermidis* AUH4567 in the TSB media (Figure S2). Under the same growth conditions (37°C, no shaking), hemin had slightly attenuated the planktonic growth at its "working" concentration of 5 µM, but it has not reduced the final OD (compared to hemin-free media). However, this slight negative impact of hemin was yet smaller compared to the impact of 5 µM N-methyl-protoporphyrin or 5 µM FeSO4.

#### *S. epidermidis* biofilm contains eDNA rich in G-quadruplex

To validate the abundance of GQs, we compared the result from two different GQ-binding antibodies: 1H6 specific to GQ-DNA (Figure S3 A, C) and BG4 specific to GQ-DNA and GQ-RNA (Figure S3 B, D). CLSM imaging revealed minor differences in the distribution of GQ-DNA identified by the two antibodies. The localization of both 1H6 and BG4 antibodies overlapped with the signal from total DNA, indicating that the antibodies did not bind to non-nucleic acid components in the biofilm. However, 1H6 bound in discrete spots while the signal from BG4 was brighter and more continuous along the web-like DNA strings. This difference may reflect the higher affinity of BG4 to GQ, or perhaps the presence of GQ-RNA in the web-like structures, as BG4 binds to both GQ-DNA and GQ-RNA while 1H6 only binds to GQ-DNA.

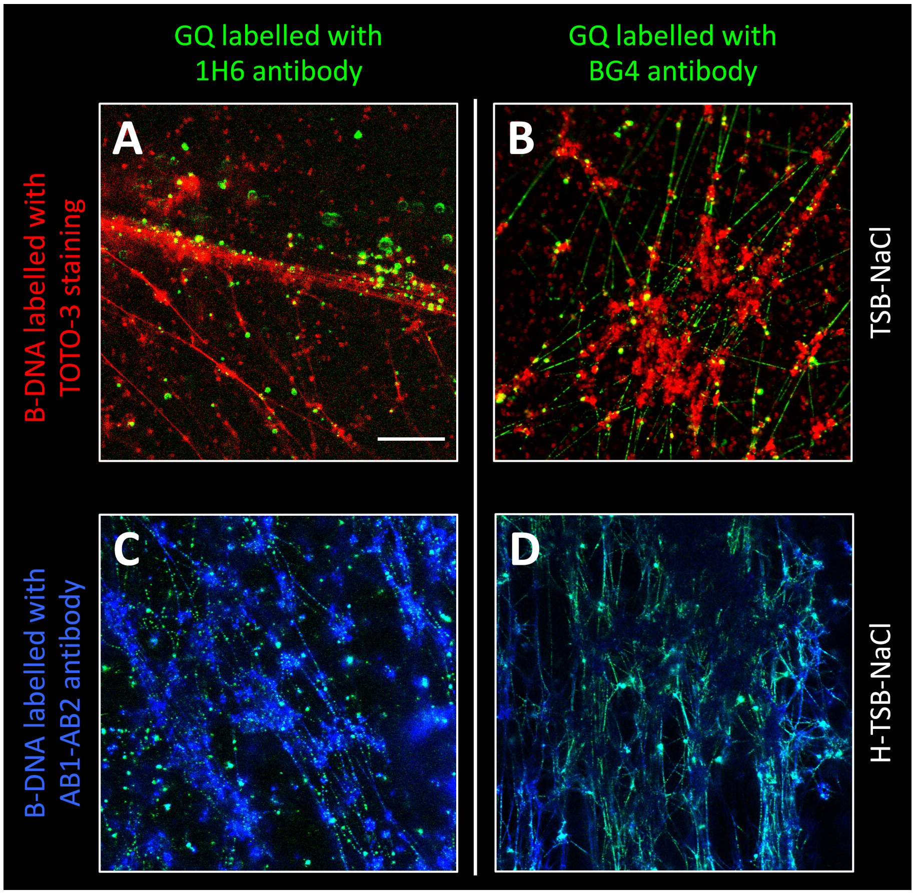


**Figure S3. Immunolabelling of GQ by 1H6 and BG4 antibodies shows stronger signal from BG4. B-DNA is best visualized by immunolabelling compared to using DNA-binding dyes**. *S. epidermidis* AUH4567 biofilms were grown for 3 days in TSB-NaCl (A, B) or H-TSB-NaCl (C, D) with 150 rpm shaking. (A, B) 2D CLSM images of GQ visualized by antibody 1H6 (A) and BG4 (B) and total eDNA visualized by TOTO-3. (C, D) 2D CLSM images of GQ visualized by antibody 1H6 (C) and BG4 (D) and total eDNA visualized by two-step AB1-AB2 immunolabelling. Scale bar 20 µm.

**B-DNA is best visualized by immunolabelling compared to using DNA-binding dyes**

To identify the most suitable method for visualizing total eDNA together with non-canonical structures, we compared the use of a DNA-binding dye (TOTO-3) to immunolabelling of B-DNA (Figure S3 A, B vs. Figure S3 C, D). It was more challenging to visualize the web-like DNA strings by TOTO-3 staining because the signal intensity from DNA strings was very low compared to other areas of the biofilm with highly concentrated eDNA. The strings could therefore easily be overlooked when imaging eDNA with DNA-binding dyes, as acquisition settings are adjusted to visualize the brightest area of the image. In contrast, the signal intensity from B-DNA visualized by immunolabelling was more homogenous (Figure S3 C, D).

Furthermore, we show comparison of the AB1-AB2 binding efficiency to the total eDNA in *S. epidermidis* AUH4567 biofilm (3-day in H-TSB-NaCl, 150 rpm) with and without prior exposure of the biofilm to 2.5 µM TOTO-3 (Figure S4). We demonstrate that TOTO-3 compromised efficiency of the eDNA immunolabelling by AB1-AB2 antibodies possibly due to its reduced affinity. We therefore chose to visualize B-DNA by immunolabelling in the subsequent experiments.

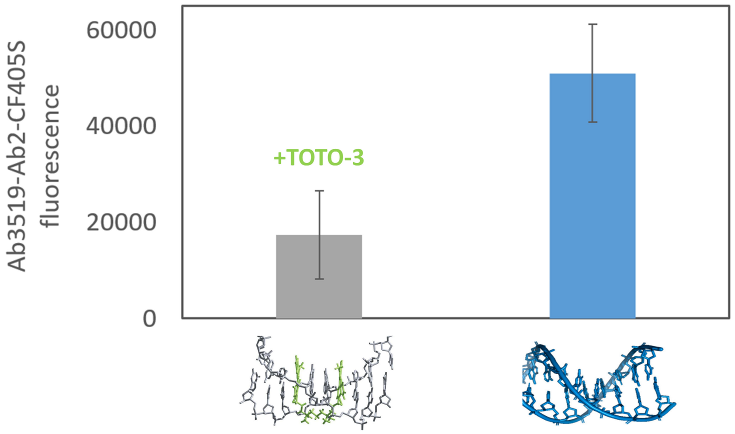


**Figure S4**. **TOTO-3 reduced efficiency of immunolabelling by AB1-AB2.** Immunolabelling of B-DNA in *S. epidermidis* AUH4567 biofilm (3-day in H-TSB-NaCl, 150 rpm) by the AB1-AB2 with and without prior exposure of the biofilm to TOTO-3. Quantification of the AB2 bulk fluorescence (n=3).

### Polysaccharides and hemin promote formation of Z-DNA in *S. epidermidis* biofilm

As a proxy-model of eDNA, we used synthetic pure DNA sequences (IDT, Table S1) annealed in the buffers optimized for GQ-DNA and Z-DNA in a thermostat.

**Table S1**. Synthetic DNA molecules used as the substrates for screening enzymes: sequences capable of folding into G-quadruplex DNA (c-myc, c-myc-4 and tel), single-stranded DNA (nonGQ, B1, B2) and their complements (B1c, B2c).

| | |
|---|---|
| **c-myc** | **GAG GGT GGG TAG GGT GGG** |
| **c-myc-4** | **GAG GGT GGG TAG GGT GGG GAG GGT GGG TAG GGT GGG GAG GGT GGG TAG GGT GGG GAG GGT GGG TAG GGT GGG CGT CAA CAG ACT CGA** |
| **Tel** | **TTA GGG TTA GGG TTA GGG TTA GGG TTA** |
| **nonGQ** | **TAG GGA TGC GAC AGA GAG GAC GGG TA** |
| **B1** | **GTG GCA GGT CAG TCA AGT ATA CTG CAC TA** |
| **B1c** | **TAG TGC AGT ATA CTT GAC TGA CCT GCC AC** |
| **B2** | **GCG CGC GCG CGC GCG CGC GCG C** |
| **B2c** | **GCG CGC GCG CGC GCG CGC GCG C** |

We used circular dichroism to identify DNA conformations formed under conditions either favoring or arresting non-canonical secondary structures. In this work, we for the first time demonstrate flipping B-DNA into Z-form by adding 0.025 % protonated chitosan (pH 5.5) to DNA solutions of B1-B1c (50 % GC) and B2-B2c (100 % GC) prior to their annealing (Figure S5 A, B). Thus, we hypothesize that the partially deacetylated PNAG in a biofilm may prompt this transition due to its polycationic nature.

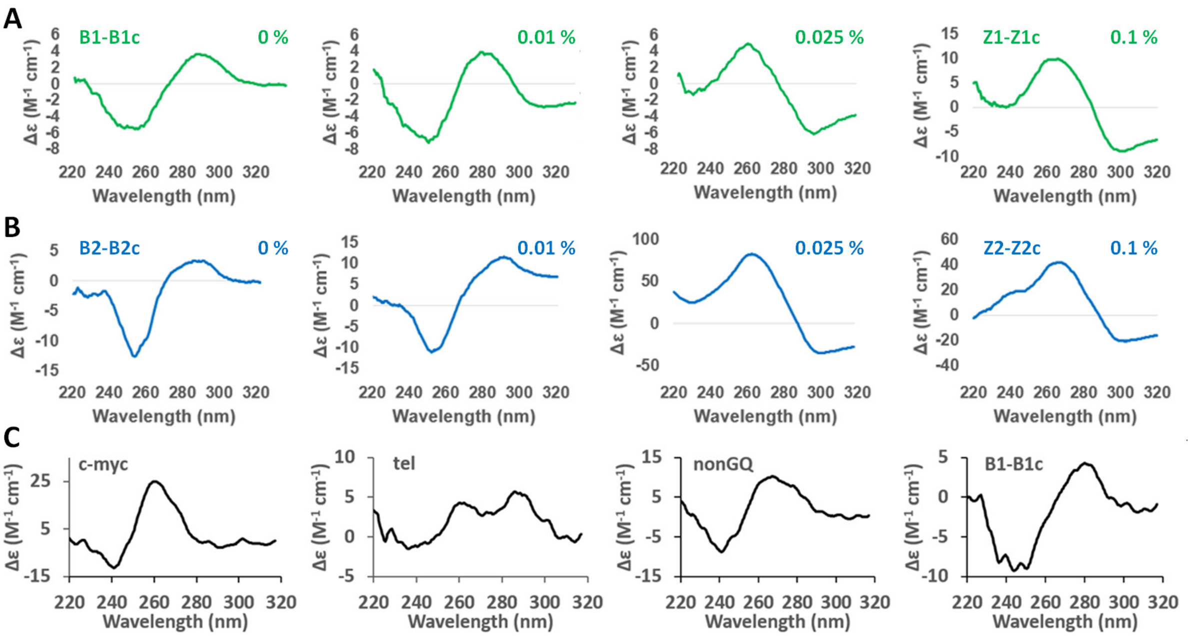


**Figure S5. DNA characterization by Circular Dichroism: chitosan flips B-DNA into Z-DNA at pH 6**. CD spectra of B-Z-DNA transition (A) between B1-B1c and Z1-Z1c and (B) between B2-B2c and Z2-Z2c as function of chitosan concentration in 25 mM Tris-acetate buffer, 6.25 mM $CaCl_2$, 1 mM $MgSO_4$ (pH 6). Flipping of B-DNA into Z-form was identified as the positive peak shift from 280 nm to 260 nm and at the same time appearance of a negative peak at 300 nm. (C) CD spectra of c-myc, tel, nonGQ and B1-B1c DNA diluted in 25 mM Tris-acetate buffer, 6.25 mM $CaCl_2$, 1 mM $MgSO_4$ (pH 6). The parallel G-quadruplex was identified as the positive peak at 260 nm.

**Hemin binds to G-quadruplex DNA in *S. epidermidis* biofilms and forms a peroxidase-like DNAzyme**

We demonstrate that GQ-DNA can be used in a biofilm as a DNAzyme with peroxidase activity. This was achieved by adding 0.1 % hydrogen peroxide and fluorescently labelled tyramide to the biofilm without adding further hemin to only detect peroxidase activity associated with hemin trapped in the biofilm. In the GQ/hemin complex, the iron center of hemin chelates hydrogen peroxide, forming an Fe(IV)-O bond and catalyzes the Fenton redox reaction, in which two electrons from a substrate (here tyramide) are transferred to hydrogen peroxide, resulting in water and a chemical bond between the tyramide and amine groups on biomacromolecules in close proximity. This process is also known as tyramide signal amplification due to the ongoing catalysis and local deposition of fluorescent tyramide near the catalyst. G-quadruplex catalyzes this chemical reaction by stabilizing intermediate complexes of the Fenton reaction through the pi-stacking.

In the biofilms that were not doped with c-myc, the peroxidase activity was also detected in non-eDNA strings, supposedly made by GQ-RNA (Figure S6).

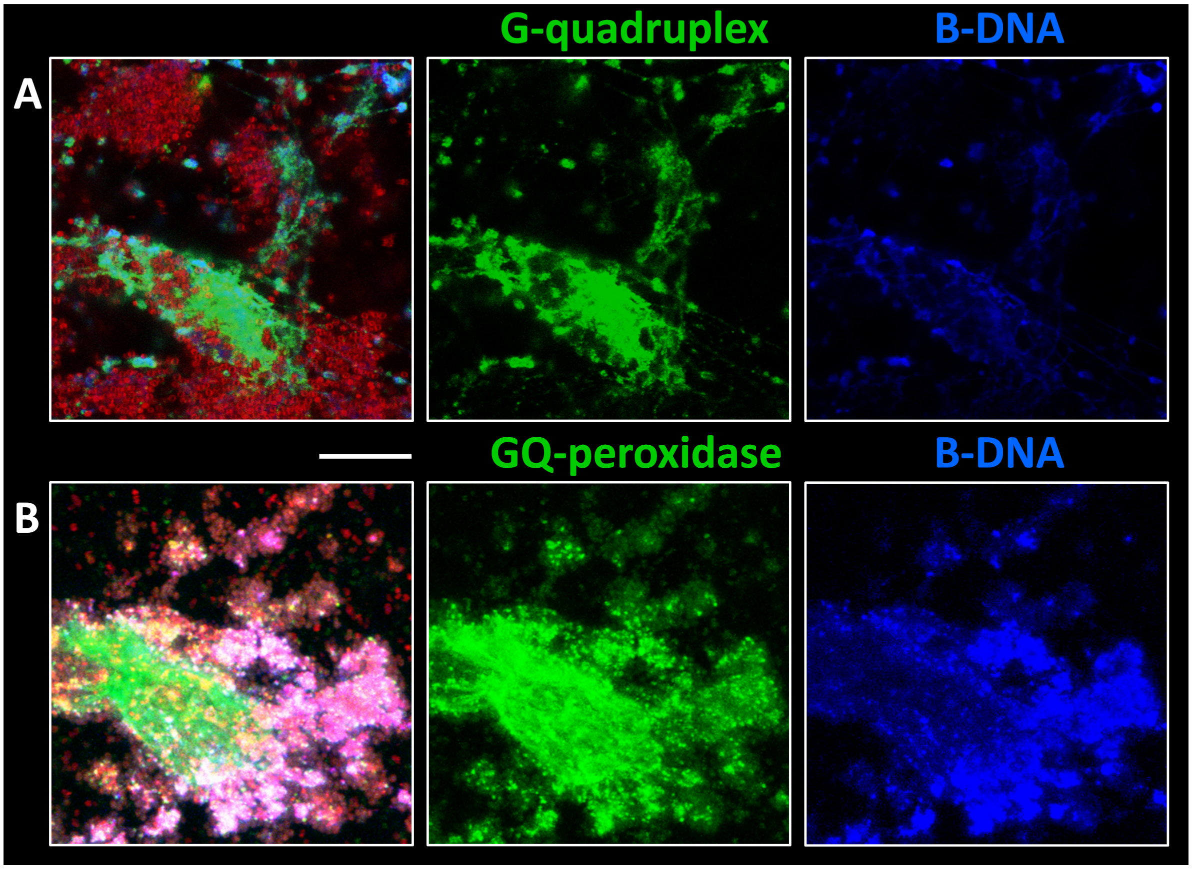


**Figure S6. Hemin and GQ-RNA enables peroxidase-like DNAzyme activity in biofilms.** S. epidermidis AUH 4567 biofilm was grown in H-TSB-NaCl for 3 days. (A) 2D CLSM image of GQ-immunolabelling by BG4 antibody (green) in combination with immunolabelling of B-DNA by AB1-AB2 antibodies (blue) and bacteria by FM 4 64 (red). (B) 3D CLSM image of peroxidase activity by tyramide labelling (green) in combination with immunolabelling of B-DNA by AB1-AB2 antibodies (blue) and bacteria by SYTO60 (red). Scale bar 20 µm.

***In vivo S. aureus* biofilm from murine osteomyelitis model contains GQ and Z-DNA**

The *in vivo* samples were visualized outside the implant area due to the implant autofluorescence in the green channel as well as little biofilm material. Here, the CLSM controls of the autofluorescence as well as further examples of the 7-day biofilms at the implant surface are presented (Figure S7).

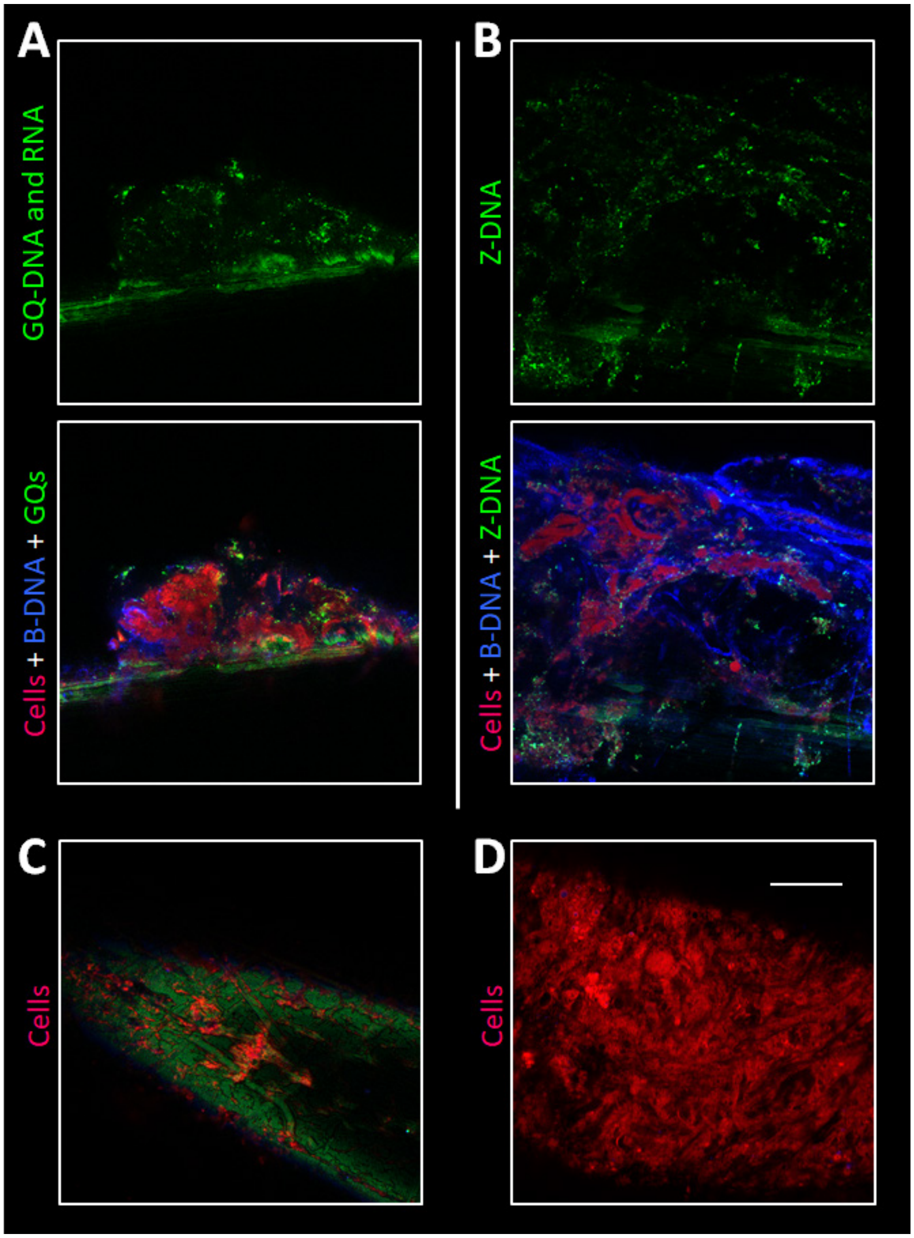


**Figure S7. In vivo biofilm from murine osteomyelitis model contains GQ-DNA and Z-DNA.** CLSM images of the tissue surrounding tibia implants infected with *S. aureus*: (A) GQ and (B) Z-DNA. (C)-(D) Samples for the autofluorescence controls were stained with FM 4 64. CLSM images were taken directly at the implant surface (C) as well as in the locations in its close vicinity (D). Cells (bacterial and murine) are shown in red (FM4-64 stain), B-DNA in blue (AB1-AB2 antibody), and GQ (BG4 antibody) or Z-DNA (Z22 antibody) in green. The 3D images are shown as single channel (green) as well as three-channel. The biofilms used for autofluorescence imaging contained only FM 4-64 stain. Scale bar 20 µm.
